# Coordinate transcriptional and post-transcriptional repression of pro-differentiation genes maintains intestinal stem cell identity

**DOI:** 10.1101/2020.06.27.175398

**Authors:** Kasun Buddika, Yi-Ting Huang, Ishara S. Ariyapala, Alex Butrum- Griffith, Sam A. Norrell, Alex M. O’Connor, Viraj K. Patel, Samuel A. Rector, Mark Slovan, Mallory Sokolowski, Yasuko Kato, Akira Nakamura, Nicholas S. Sokol

## Abstract

The role of Processing bodies (P-bodies), key sites of post-transcriptional control, in adult stem cells remains poorly understood. Here, we report that adult Drosophila intestinal stem cells, but not surrounding differentiated cells such as absorptive Enterocytes (ECs), harbor P-bodies that contain *Drosophila* orthologs of mammalian P-body components DDX6, EDC3, EDC4 and LSM14A/B. A targeted RNAi screen in intestinal progenitor cells identified 39 previously known and 64 novel P-body regulators, including *Patr-1*, a gene necessary for P-body assembly. Loss of *Patr-1*-dependent P-bodies leads to a loss of stem cells that is associated with inappropriate translation and expression of EC-fate gene *nubbin*. Transcriptomic analysis of progenitor cells identifies a cadre of such weakly transcribed pro-differentiation transcripts that are elevated after P-body loss. Altogether, this study identifies a coordinated P-body dependent, translational and transcriptional repression program that maintains a defined set of *in vivo* stem cells in a state primed for differentiation.

**Graphical abstract:** 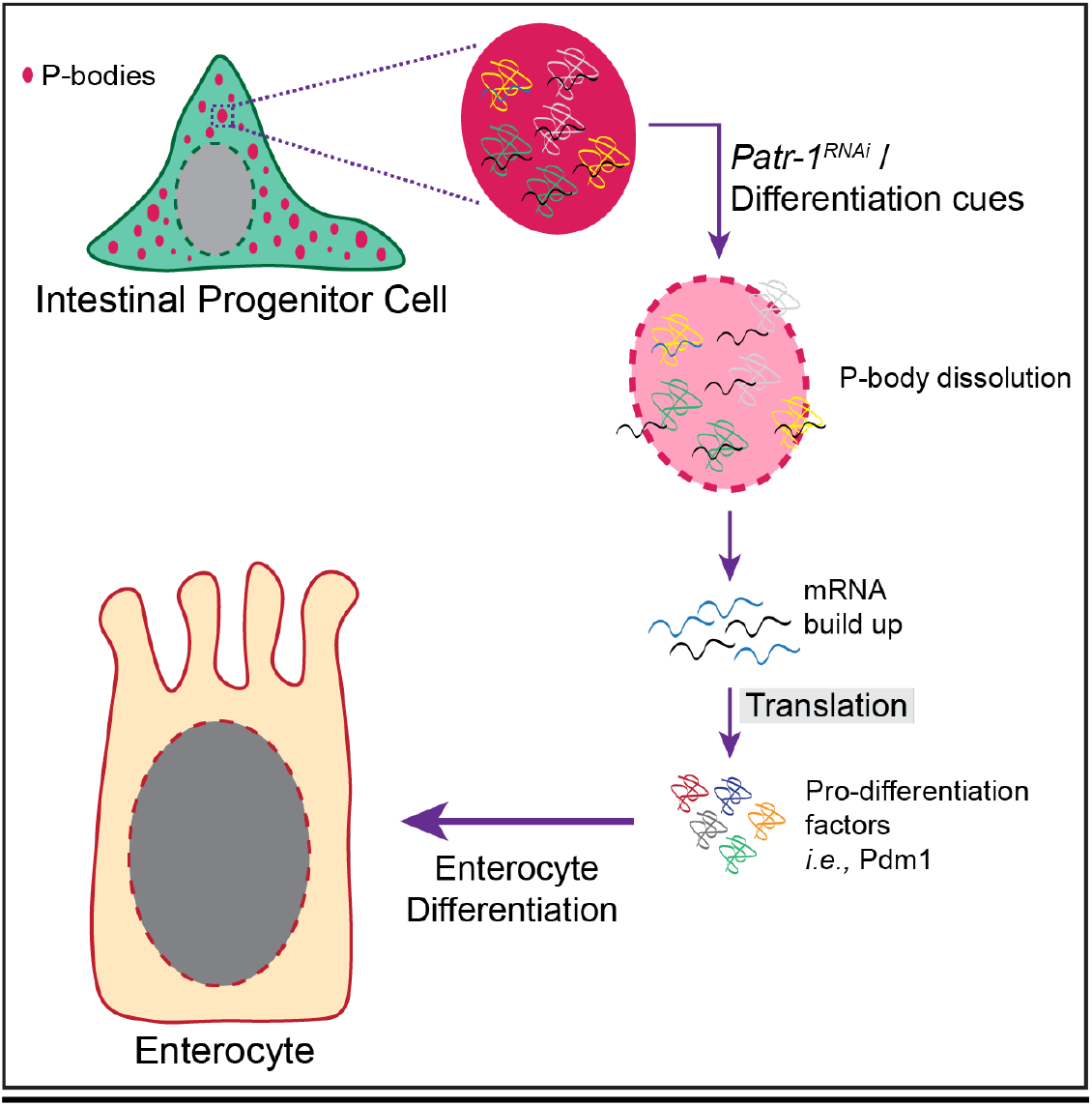

**Highlights:**

- Drosophila intestinal progenitor cells contain constitutive and ultrastructurally organized P-bodies.
- A P-body regulator *Patr-1* is required for intestinal progenitor cell maintenance.
- Enterocyte (EC) genes such as *nubbin* are weakly transcribed but not translated in intestinal progenitors.
- P-bodies repress EC gene translation to promote stem cell maintenance.

## Introduction

In eukaryotic cells, translationally inactive messenger RNAs (mRNAs) can assemble to form a distinct population of cytoplasmic messenger ribonucleoprotein (mRNP) particles known as Processing bodies (P-bodies) (Anderson and Kedersha, 2006; Decker and Parker, 2012; Kedersha et al., 2005). Like stress granules, P-bodies lack a limiting membrane and are visible via conventional microscopic techniques (Anderson and Kedersha, 2006), are enhanced or induced by blocking translation initiation (Buchan and Parker, 2009; Chan et al., 2018; Franks and Lykke-Andersen, 2008; Kedersha et al., 2000; Teixeira et al., 2005), are disassembled when treated with cycloheximide, a chemical that traps mRNAs on ribosomes (Anderson and Kedersha, 2009; Buchan and Parker, 2009; Kedersha et al., 2000; Mugler et al., 2016; Teixeira et al., 2005), and are conserved from yeast to humans (Martínez et al., 2013; Protter and Parker, 2016). Unlike stress granules, however, P-bodies are constitutive and contain proteins involved in mRNA decay (Anderson and Kedersha, 2006, 2009; Buchan and Parker, 2009; Decker and Parker, 2012; Eulalio et al., 2007a; Luo et al., 2018; Parker and Sheth, 2007; Sheth and Parker, 2003; Youn et al., 2019). While recent studies have identified stress granules in stem cells of adult tissues (Buddika et al., 2020a), whether P-bodies are also present in adult stem cells and, if so, what biological processes they might control is not well understood.

Recent biochemical profiling experiments in human HEK293 cells identified different classes of P-body resident proteins that include translation repression and mRNA decay factors (4E-T, LSM14A, LSM14B and IGF2BP2), microRNA pathway components (AGO1, AGO2 and MOV10), decapping complex proteins (DCP1A, DCP1B, DCP2 and EDC4) and nonsense-mediated mRNA decay (NMD) factors (UPF1) (Hubstenberger et al., 2017). Similar studies have also identified 6,168 transcripts to be significantly enriched in P-bodies and another 7,588 transcripts to be significantly depleted (Hubstenberger et al., 2017). This clear bifurcation of mRNAs indicates that the P-body transcriptome is large but specific (Hubstenberger et al., 2017; Standart and Weil, 2018). While its protein composition suggests that the primary function of P-bodies is mRNA degradation (Anderson and Kedersha, 2006; Decker and Parker, 2012), recent studies have found that mRNAs localized to P-bodies can return to the cytoplasm to re-enter translation (Bhattacharyya et al., 2006; Brengues et al., 2005) and that P-bodies can be disrupted without affecting mRNA decay (Eulalio et al., 2007a; Stoecklin et al., 2006). Therefore, current models propose that P-bodies coordinate the storage of mRNA regulons leading to stoichiometric protein production (Standart and Weil, 2018). However, genetically tractable *in vivo* P-body models are necessary to better understand the exact role of P-bodies in eukaryotic tissues including the identification and tracking of P-body regulated mRNAs *in vivo*. Developing such tissue-based models will also provide insights into the structural organization of P-bodies. While electron micrographs of P-bodies in HeLa cells have identified densely populated structures within P-bodies (Souquere et al., 2009), the integration of super-resolution based characterizations are required to elucidate whether P-body resident components are structurally organized (Youn et al., 2019).

Although some post-transcriptional processes such as alternative splicing, polyadenylation, and RNA modifications have been extensively studied in pluripotent stem cells (Chen and Hu, 2017), the roles of these and other RNA processes, such as RNA decay, storage, and translational control, in the potency of adult somatic stem cells remain poorly understood. For example, while recent work indicated that post-transcriptional regulation of mRNAs via P-bodies play vital roles in adult stem cell proliferation and differentiation, it is complex and dynamic: functional analysis of the P-body protein DDX6 found that it promotes differentiation in some *in vitro* derived stem cell lineages and represses it in others (Di Stefano et al., 2019). Thus, tissue-based stem cell models will be critical for probing P-body function in an *in vivo* context. The intestinal epithelium of adult *Drosophila* offers such a model to study post-transcriptional gene regulatory mechanisms (Buddika et al., 2020a; Chen et al., 2015; Luhur et al., 2017). This actively regenerating tissue hosts a population of progenitor cells, which is composed of two main cell types: mitotic intestinal stem cells (ISCs) and their transient and non-mitotic daughters, enteroblasts (EBs) (Micchelli and Perrimon, 2006; Ohlstein and Spradling, 2006). In this study we used the Drosophila midgut as a tissue-based stem cell model to delineate the biological roles of P-bodies *in vivo*.

## Results

### P-body components are enriched in intestinal progenitor cells

To test whether intestinal progenitor cells contained P-bodies, we stained intestines dissected from 7-day old, adult female *Drosophila* with a series of four antibodies. These detected the Drosophila orthologs of known P-body proteins in other systems: EDC3, GE-1 (known in metazoans as EDC4), Me31B (known in metazoans as DDX6), and TRAL (known in metazoans as LSM14A) (Anderson and Kedersha, 2006; Ayache et al., 2015; Eulalio et al., 2007a; Hubstenberger et al., 2017). Using previously verified antibodies for GE-1 and Me31B (Barbee et al., 2006; Fan et al., 2011; Nakamura A, Amikura R, Hanyu K, 2001) as well as new, validated antibodies for TRAL and EDC3 (**Fig. S1A-B**), we found that all these antibodies displayed elevated signal in intestinal progenitors (**Fig. 1A-D**), which were labeled by horseradish peroxidase (HRP) (Miller et al., 2020; O’Brien et al., 2011). Limited to the cytoplasm, confocal microscopy detected these proteins in clearly separated cytoplasmic granules that were similar in level and organization in both ISCs and *3Xgbe-smGFP::V5::nls* positive EBs (**Fig. 1A’-D’**). TRAL and Me31B staining perfectly overlapped, indicating colocalization (**Fig. 1E-E’**). The relative absence of staining in neighboring polyploid ECs also indicated that these proteins were downregulated during EC differentiation (see **Fig. 1A-E**). To assess whether such downregulation occurred during EE differentiation as well, we co-stained intestines for Prospero (PROS), an EE-marker (Micchelli and Perrimon, 2006), and either TRAL or Me31B. While the majority of PROS+ cells showed very low to no detectable TRAL or Me31B, ~20% displayed levels that were comparable to progenitor cells (**Fig. S1C-D**). These data indicated the presence of a population of cytoplasmic granules containing colocalized P-body components in progenitor cells that were cleared during EC and EE differentiation.

**Figure 1:**
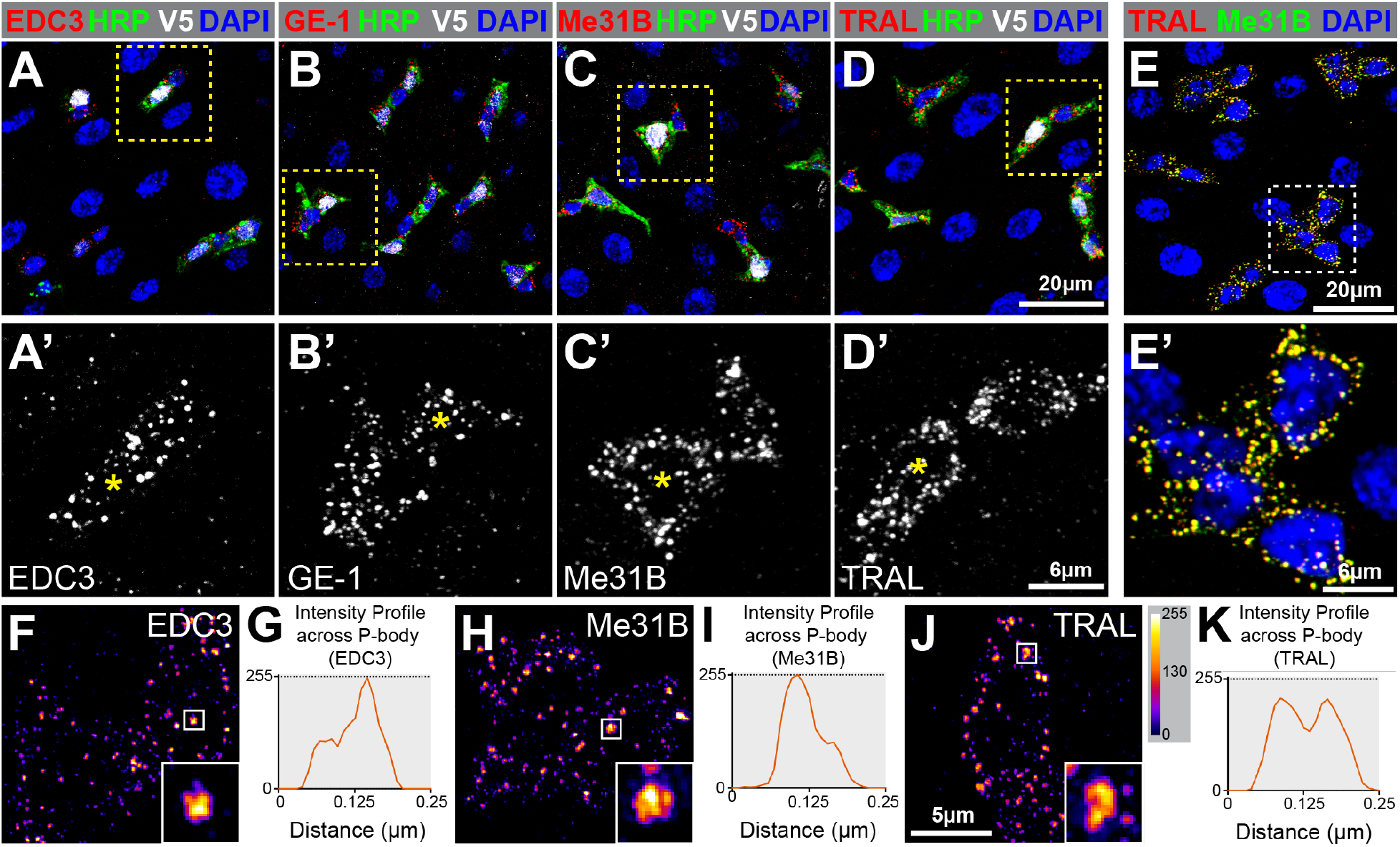
Intestinal progenitor cells contain P-bodies. (A-D) Adult posterior midguts stained in red for (A) EDC3, (B) GE-1, (C) Me31B, and (D) TRAL. *3Xgbe-smGFP::V5::nls* (white) labels EBs, HRP (green) labels both EBs and ISCs, and the DNA-dye DAPI (blue) labels all cells. A’-D’ are enlargements of progenitor cells indicated in A-D with EBs labeled (asterisks). (E) Adult posterior midgut stained for TRAL (red), Me31B (green), and DAPI (blue). (F, H, J) Super-resolution micrographs of progenitor cells stained for (F) EDC3, (H) Me31B, or (J) TRAL and pseudocolored using scale in J to indicate protein level. Insets are enlarged views of boxed regions. (G, I, K) Line plots showing pixel-by-pixel intensity profiles of insets.

### Intestinal P-bodies are ultrastructurally organized

Because the structural organization of P-bodies is not well understood (Youn et al., 2019), we used this system to assess the ultrastructural organization of intestinal P-body protein complexes. We stained dissected intestines with either Me31B, TRAL or EDC3, imaged the cytoplasmic distribution of these proteins using structured illumination super-resolution microscopy (SIM), and then used the FIJI package of ImageJ to generate intensity maps of each protein. These maps revealed that the distribution of Me31B, TRAL, and EDC3 was not uniform within a granule (**Fig. 1F, H, J**). We noted that the pixel density was different along the length of a granule and more central domains within the granule contained greater protein accumulation than the rest of the granule (**Fig. 1G, I, K**). Given that this non-uniform protein organization within mature P-bodies is very similar to the core-shell organization of a stress granule, we propose that P-bodies share the same structural organization. Although significantly smaller, P-bodies are likely to have multiple cores similar to stress granules. Taken together, these data identified the existence of a structurally organized population of steady-state P-bodies in intestinal progenitors of adult *Drosophila*.

### P-bodies and stress granules are distinct mRNPs in progenitor cells

The above analysis suggested that P-body complexes were distinct from intestinal progenitor stress granules (IPSGs) because P-bodies were present under standard conditions while IPSG complexes only formed after intestines were exposed to acute stress (Buddika et al., 2020a). We therefore compared the subcellular locations and sizes of P-body proteins (TRAL/Me31b) with IPSG proteins (Fragile Mental Retardation Protein [FMRP]/Rasputin[RIN]) in both unstressed and stressed conditions using SIM. For these experiments, unstressed samples were prepared by incubating dissected intestines *ex-vivo* in Krebs Ringer Bicarbonate Buffer (KRB) for 60 minutes; stressed intestines were treated with 1mM rapamycin in KRB for the same length of time. P-body proteins perfectly colocalized with each other under unstressed conditions but only partially colocalized with IPSG proteins under these same conditions (**Fig. S1E-J**). However, after stress, P-body and IPSG proteins perfectly localized (**Fig. S1K-P**). To investigate the sizes of these complexes, we measured the sizes of TRAL and FMRP punctae. Under unstressed conditions, TRAL punctae (0.047 µm^2^; n=804 puncta in 14 cells) were bigger than FMRP punctae (0.026 µm^2^; n=629 puncta in 8 cells] (**Fig. S1Q-R, U**). Stress treatment caused TRAL punctae (0.097 µm^2^; n=657 puncta in 25 cells) and FMRP punctae (0.098 µm^2^; n=1215 puncta in 42 cells) to grow to the same size (**Fig. S1S-U**). From this analysis, we concluded that P-body proteins localized to persistent mRNP complexes that, unlike IPSGs, were present in unstressed cells, and that stress led to the enlargement of these mRNPs into IPSGs.

### Identification of genetic modifiers of progenitor P-bodies using a targeted screen

To characterize the genetic framework governing P-body formation in progenitor cells, we performed a targeted RNAi screen to identify genes that altered P-body morphology. We focused on genes implicated in RNA-related processes, and generated a list of 600 candidate genes by using the GO term analysis function on Flybase. We selected the 485 genes with Bloomington *Drosophila* Stock Center RNAi strains to conduct a primary screen, and used TRAL as the P-body marker to score for P-body morphology (**Table S1**). We screened a total of 735 RNAi strains that targeted these 485 genes: 186 genes were targeted using two or more RNAi strains, and the remainder were targeted by just one RNAi line. Individual RNAi strains were crossed to a conditional version of the progenitor cell driver *esg-GAL4* (*esgTS*) (Micchelli and Perrimon, 2006), induced at the adult stage, and dissected seven days later. From this analysis, 103 genes that affected TRAL pattern or level were identified and grouped into five phenotypic classes: [1] large TRAL puncta (“Big”), [2] small TRAL puncta (“Small”), [3] dispersed TRAL puncta (“Diffuse”), [4] both dispersed and large TRAL puncta (“Diffuse + Big”), and [5] no TRAL staining (“No TRAL”) (**Fig. 2A-B**). To assess the specificity of these effects on P-body morphology, a representative set of 25 positives -- 13 “big” genes, 3 “small” genes, 4 “diffuse” genes, 3 “diffuse + big” genes, and 2 “no TRAL” genes -- were also co-stained with IPSG-makers FMRP and ROX8 (**Fig. S2A-F, Table S1**). While most had no discernable effect, two genes, *fne* and CG13928, altered the expression profile of these IPSG proteins and led to “big” or “diffuse and big” FMRP/ROX8 foci, respectively (**Table S1, Fig. S2E**). As expected, many of the positive genes encode proteins that were implicated in mRNA metabolic processes (54 genes) and were known components of ribonucleoprotein complexes (42 genes) (**Fig. S2G-H**). For example, the targeting of genes encoding P-body resident proteins EDC3 or PCM, the *Drosophila* homolog of mammalian 5’-3’ exonuclease XRN1, led to bigger TRAL granules. While 39 (38%) of the identified genes were orthologs of proteins known to affect P-bodies in human cells (**Table S1**) (Youn et al., 2019), we also identified 64 (62%) genes that were not previously implicated in P-body organization or assembly. Unexpectedly, regulators of nuclear processes such as RNA splicing (35 genes) and nuclear export (9 genes) as well as components of nuclear assemblies such as the exon-exon junction complex (6 genes) also impacted the morphology of cytoplasmic TRAL granules (**Fig. S2G-H, Table S1**). We also noted that knockdown of 24% of the positive genes (23/103 genes), caused loss of progenitor cells. These results therefore raise the possibility that P-bodies are physiologically linked to the stemness of this progenitor cell population.

**Figure 2:**
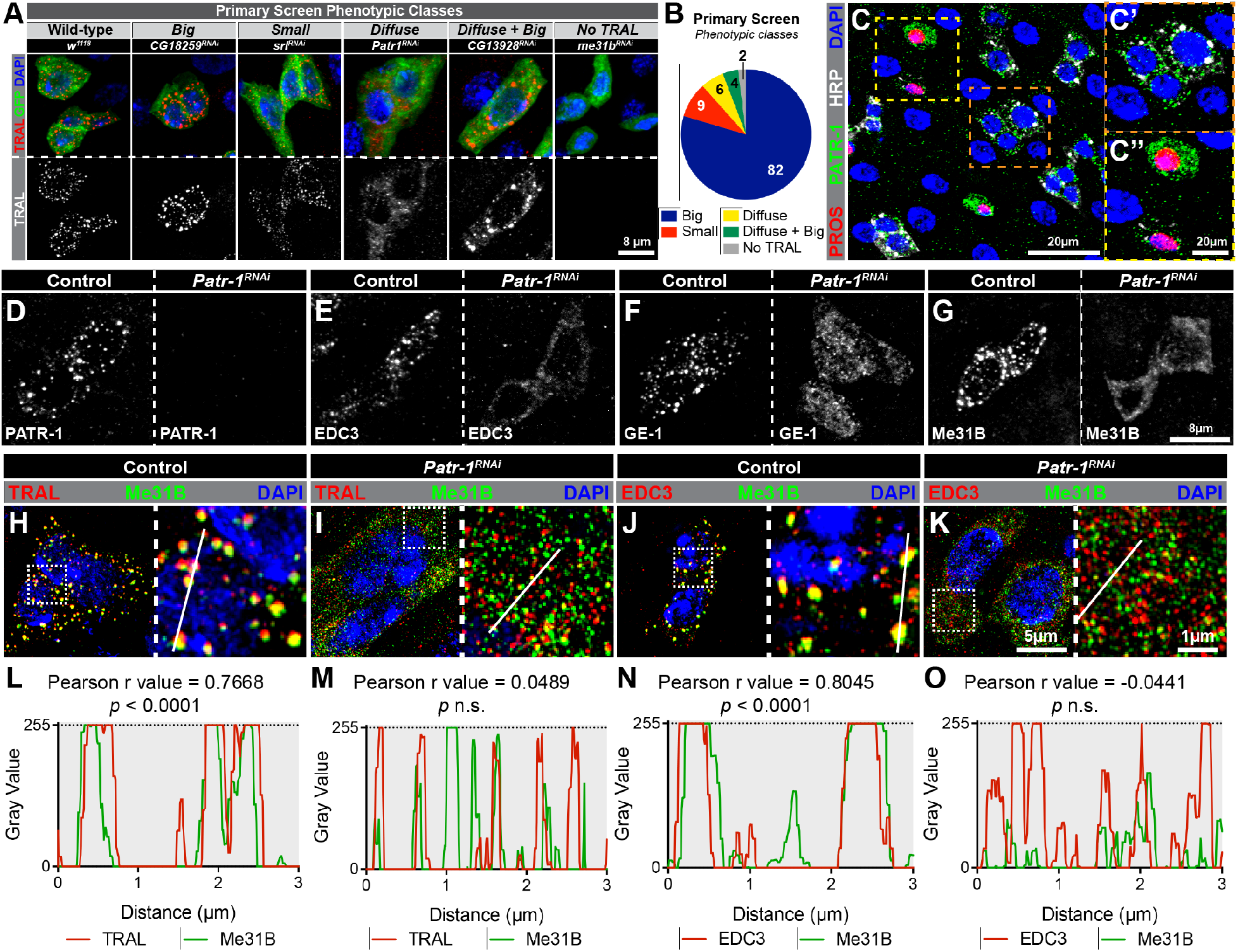
RNAi screen identifies PATR-1 as an essential component of progenitor cell P-bodies. (A) Examples of the five classes of P-body phenotypes showing progenitor cells from *esg*^*TS*^, *esg*^*TS*^ / *CG18259* RNAi, *esg*^*TS*^ / *srl* RNAi, *esg*^*TS*^ / *Patr-1* RNAi, *esg*^*TS*^ / *CG13928* RNAi, or *esg*^*TS*^ / *me31b* RNAi intestines stained for TRAL (red), GFP (green), and DAPI (blue). (B) Chart of phenotypic class numbers. (C) Posterior midguts stained for PROS (red), PATR-1 (green), HRP (white) and DAPI (blue). C’ and C’’ are enlargements of regions indicated in C. (D-G) Progenitor cells from *esg*^*TS*^ (control, left) or *esg*^*TS*^ */ Patr-1* RNAi (right) intestines stained for (D) PATR-1, (E) EDC3, (F) GE-1, or (G) Me31B. (H-K) Super-resolution micrographs of (H, J) *esg*^*TS*^ or (I, K) *esg*^*TS*^ */ Patr-1* RNAi intestinal progenitors stained for Me31B (green), DAPI (blue) and either TRAL (red in H, I) or EDC3 (red in J, K). (L-O) Normalized pixel-by-pixel fluorescent intensity profiles of indicated protein (TRAL, Me31B or EDC3) along white lines in H-K.

### PATR-1 is a P-body resident protein that is necessary for their formation

To investigate the function of P-bodies in progenitor cells in more detail, we focused on *Patr-1*, a member of the “diffuse” class and a known critical nucleator of P-bodies in other systems (Coller and Parker, 2005; Nissan et al., 2010; Pilkington and Parker, 2008; Pradhan et al., 2012; Sachdev et al., 2019; V et al., 2011). Using a verified antibody (Pradhan et al., 2012), we found PATR-1, like other mRNP proteins, was almost exclusively detected in the cytoplasm of progenitor cells and displayed a distinctive granular organization (**Fig. 2C, 2C’**). PATR-1 also showed a strong but less punctate expression in ~45% of EEs (compare **Fig. S1C-D** to **Fig. 2C’’**). PATR-1 colocalized well with both TRAL and Me31B (**Fig. S2I-J, L-M**) and did not colocalize with the IPSG protein FMRP (**Fig. S2K, N**), indicating it was a P-body component. We therefore used the PATR-1 antibody as an independent marker to verify the effects of the 103 genes identified above on P-body morphology and found analogous effects in all cases except for the “No TRAL” category (**Fig. S2O-T**). In this class, smaller than normal PATR-1 granules were observed (**Fig. S2T**), suggesting that the identified genes altered overall P-body morphology rather than specifically just TRAL staining. Collectively, these data indicated that PATR-1 was a P-body component in intestinal progenitor cells of adult *Drosophila*.

To test whether PATR-1 was required for the recruitment of P-body proteins in addition to TRAL, we first verified that *Patr-1* RNAi eliminated PATR-1 protein expression (**Fig. 2D**) and then analyzed the effect of *Patr-1* RNAi on EDC3, GE-1 and Me31B distribution. The prominent punctate pattern and colocalization of these proteins was lost after *Patr-1* RNAi (**Fig. 2E-O**) indicating a failure in P-body assembly as well as P-body protein interaction. To verify these RNAi phenotypes, we generated a new series of *Patr-1* null alleles that were homozygous lethal, died as third instar larvae, displayed the same lethal phase when in *trans* to a deficiency, and lacked PATR-1 protein by Western blot (**Fig. S3A-B**). Using two representative alleles, *Patr-1*^*P105Δ*^ and *Patr-1*^*R107FS1*^, we then generated *Patr-1* mutant adult progenitors with the Mosaic Analysis with a Repressible Cell Marker (MARCM) technique (Lee and Luo, 1999).

Seven days after clone induction, intestines were dissected and either stained for PATR-1 or scored for P-body morphology based on Me31B and EDC3 distribution. PATR-1 staining was not detected in either *Patr-1*^*P105Δ*^ or *Patr-1*^*R107FS1*^ homozygous mutant cells (**Fig. S3C-D**) and a previously verified rescue transgene, *P21M20* (Pradhan et al., 2012), restored PATR-1 staining to *Patr-1*^*P105Δ*^ mutant cells (**Fig. S3E**). Similar to *Patr-1* RNAi, the P-body localization of Me31B and EDC3 was completely disrupted in *Patr-1*^*P105Δ*^ and *Patr-1*^*R107FS1*^ mutant clones (**Fig. S3F-G** and **I-J**), and these defects were rescued entirely by the *P21M20* transgene (**Fig. S3H** and **K**). Taken together, these data indicated that PATR-1 was required for P-body assembly in the cytoplasm of adult *Drosophila* intestinal progenitor cells.

### Loss of PATR-1 leads to progenitor cell loss

To analyze the effects of P-body loss on progenitor cell biology, we crossed control and *Patr-1* RNAi strains to *esgTS* and analyzed intestines for an effect on the total number of cells as well as the number of either progenitors, EEs, or ECs at three timepoints (**Fig. 3A**). Intestinal progenitor cell number in *Patr-1* RNAi intestines gradually declined over time and was almost completely absent after 15 days of RNAi (**Fig. 3B,F-I**). In contrast, EE cell percentage remained unchanged at early time points and showed a slight increase at 15 days (**Fig. 3C**). In addition, EC percentage increased over time and showed ~30% increase at 15 days (**Fig. 3D**). However, the total number of cells per field was significantly reduced at 15 days (**Fig. 3E**), at least in part reflecting the loss of progenitor cells. Consistent with these results, *Patr-1*^*P105Δ*^ and *Patr-1*^*R107FS1*^ mutant clones were smaller and contained fewer ISCs compared to control (**Fig. S3L-R**); these defects were rescued entirely by the *P21M20* transgene. To further validate the loss of mitotic cells in the absence of PATR-1, we fed control and flies expressing *Patr-1* RNAi (14 days) for 1 day with bleomycin, a chemical that induces rapid ISC proliferation due to EC damage, and quantified mitotic proliferation by staining for phosphorylated-histone 3 (pH3), a highly specific marker for condensed chromosomes during mitosis (Amcheslavsky et al., 2009). Consistent with progenitor cell loss, intestines expressing *Patr-1* RNAi for 15 days displayed significantly reduced numbers of pH3 positive cells (**Fig. S3S**). Taken together, these data suggested that progenitor cells were lost and the epithelial integrity was compromised due to a failure in P-body assembly.

**Figure 3:**
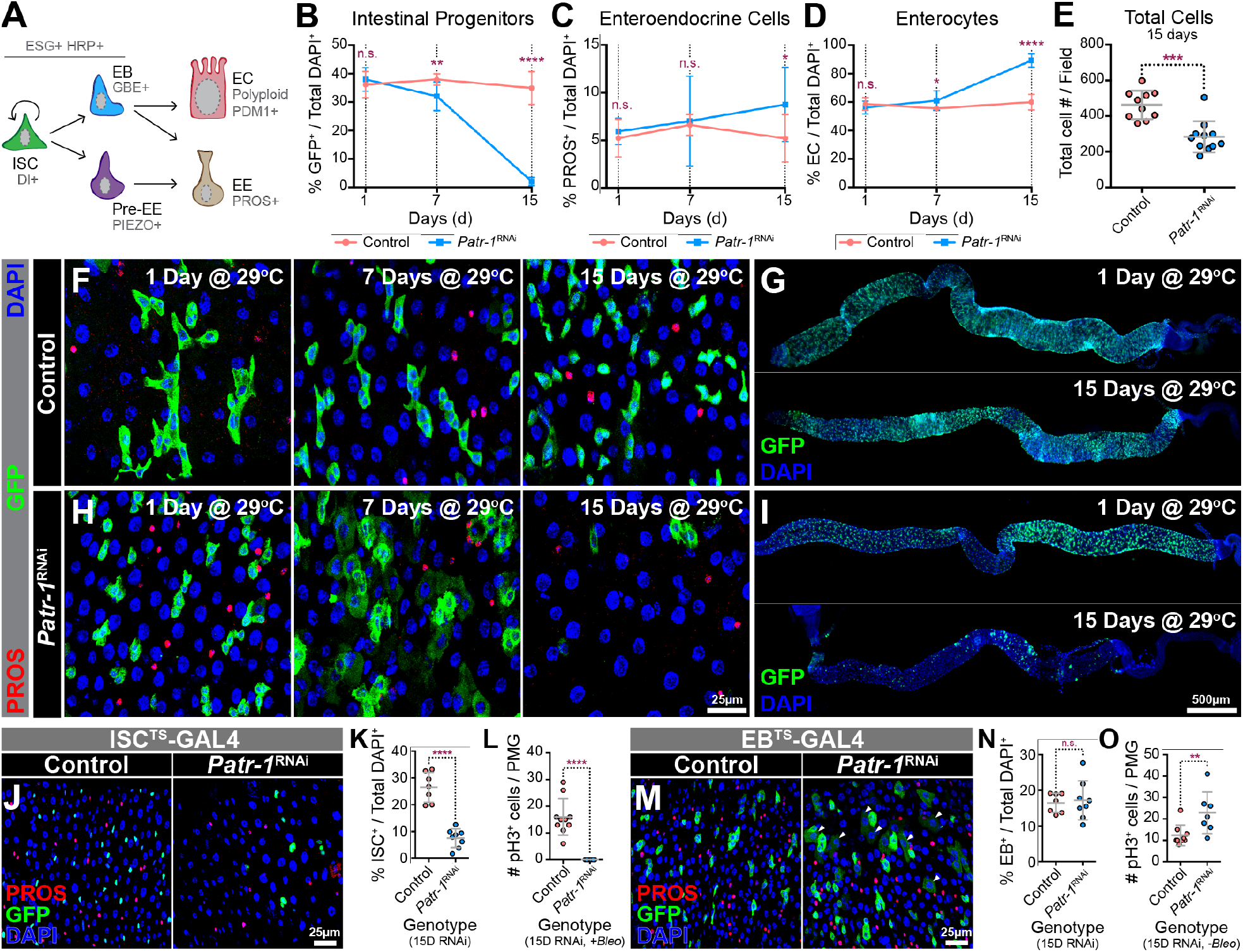
PATR-1 maintains intestinal progenitor cells. (A) Schematic of intestinal cell types and markers. (B-D) Normalized (B) intestinal progenitor cell, (C) EE cell, or (D) EC percentages in *esg*^*TS*^ (pink) and *esg*^*TS*^ */ Patr-1* RNAi (blue) posterior midguts (n=8-to-11) after 1, 7 and 15 days at 29°C. (E) Dot-plot of total cell number per field of view in *esg*^*TS*^ (n=10) and *esg*^*TS*^ */ Patr-1* (n=11) midguts after 15 days at 29°C. (F, H) Sections from (F) *esg*^*TS*^ or (H) *esg*^*TS*^ */ Patr-1* RNAi midguts after 1, 7 and 15 days at 29°C stained for PROS (red), GFP (green) and DAPI (blue). (G, I) *esg*^*TS*^ (G) and *esg*^*TS*^ */ Patr-1* RNAi (I) intestines after 1 and 15 days at 29°C stained for GFP (green) and DAPI (blue). (J) Sections from *ISC-KCKT-GAL4*^*TS*^ (left) or *ISC-KCKT-GAL4*^*TS*^ */ Patr-1* RNAi (right) midguts after 15 days at 29°C stained for PROS (red), GFP (green) and DAPI (blue). (K-L) Normalized ISC percentage (n=7 or 8) or number of pH3+ cells per posterior midgut after bleomycin feeding (n=10 or 7) of genotypes shown in J. (M) Sections from *gbe*^*TS*^ (left) or *gbe*^*TS*^ */ Patr-1* RNAi (right) 15 days after shifting to 29°C and stained for PROS (red), GFP (green) and DAPI (blue). (N-O) Normalized EB percentage (n=7 or 8) or the total number of pH3+ mitotic cells per posterior midgut (n=9 or 7) without bleomycin feeding of genotypes shown in M. Error bars on plots show mean±s.d. and asterisks denote statistical significance from Unpaired t-test (B-D, K, N-O) or Mann-Whitney test (E, L). \**p* <0.05; \*\**p* < 0.01; \*\*\**p* < 0,001; \*\*\*\**p* < 0.0001; n.s., not significant.

Given that P-bodies were present in both ISCs and EBs, we assessed whether P-body function was necessary in each cell type by performing knockdown of *Patr-1* using conditional, cell type specific GAL4 drivers, denoted as *ISC*^*TS*^*-* and *EB*^*TS*^*-GAL4* respectively (Buddika et al., 2020b; Zeng et al., 2010). Consistent with *esg-GAL4* based knockdown, knocking down *Patr-1* in ISCs for 15 days led to a marked reduction in GFP positive ISCs (**Fig. 3J-K)** as well as the complete absence of pH3 positive cells after bleomycin feeding (**Fig. 3L**). Thus, the few surviving ISCs had lost the proliferative potential that wild type ISCs possess. In contrast to the requirement of *Patr-1* for ISC maintenance, knocking down *Patr-1* using the *EB*^*TS*^*-GAL4* had no notable effect on GFP positive EB cell number compared to control after 15 days at permissive temperature (**Fig. 3M-N**). However, *Patr-1* knockdown in EBs resulted in both a marked increase in ISC proliferation (**Fig. 3O**) as well as the emergence of a weakly GFP positive EC-like polyploid cell population (**Fig. 3M**, right). These observations suggested that *Patr-1* loss caused both an increase in the rate of EB differentiation and a commensurate increase in the rate of ISC proliferation that maintained EB number at normal levels. Taken together, these results indicated that *Patr-1* was required to maintain ISCs and limit the rate of EB differentiation.

### PATR-1 represses pro-differentiation genes in intestinal progenitors

To identify P-body dependent mRNAs whose misregulation might be the underlying cause of the progenitor cell loss described above, we performed transcriptomic profiling (RNA-seq) of FACS (fluorescent activated cell sorting) isolated *Patr-1* RNAi and control intestinal progenitor cells (**Fig. 4A, S4A**). *Patr-1* knockdown in intestinal progenitors significantly altered the expression of 1,226 transcripts (**Fig. 4B, Table S2**); 703 were upregulated and 523 were downregulated including, most significantly, *Patr-1* mRNA (**Fig. 4B-D, Table S2**). Because P-body regulated mRNAs tend to be longer and have a lower GC content than average (Courel et al., 2019), we evaluated the misregulated genes for these characteristics: upregulated genes (3336.1 ± 143.9 bp; 44.1 ± 0.2 %) were significantly longer and had a lower GC content than unchanged genes (2698.4 ± 27.7 bp; 46.1 ± 0.1 %) (**Fig. S4B**). Gene Ontology (GO) analysis showed that cell differentiation genes were significantly upregulated, in agreement with the identities of mRNAs found in HEK293 P-bodies (Hubstenberger et al., 2017) (**Fig. S4C**). Since the expression of key pro-apoptotic and -EE fate mRNAs were unchanged (**Fig. S4D-E, Table S2**), this transcriptomic analysis suggested that *Patr-1* mutant progenitor cells were lost due to elevated differentiation caused by inappropriate expression of pro-EC fate genes. Similar effects on EC genes have previously been associated with phenotypes caused by loss of other progenitor genes, including the Snail family transcription factor Escargot (Esg) and its co-factor Verthandi (Vtd or Rad21) (Khaminets et al., 2020; Korzelius et al., 2014). Notably, 22% (92/410 genes) and 18% (80/454 genes) of the significantly upregulated genes in *esg*-RNAi and *vtd*-RNAi overlapped with *Patr-1* RNAi upregulated genes (**Fig. S4F-G, Table S2**). These included EC-marker genes such as *nubbin* (*nub*, also known as POU-domain protein 1 or *Pdm1*), *Myo61F*, and *big bang* (*bbg*) (Khaminets et al., 2020; Korzelius et al., 2014). Taken together, these results suggested that Esg/Vtd and *Patr-1* transcriptionally and post-transcriptionally repress an overlapping cohort of pro-EC genes.

**Figure 4:**
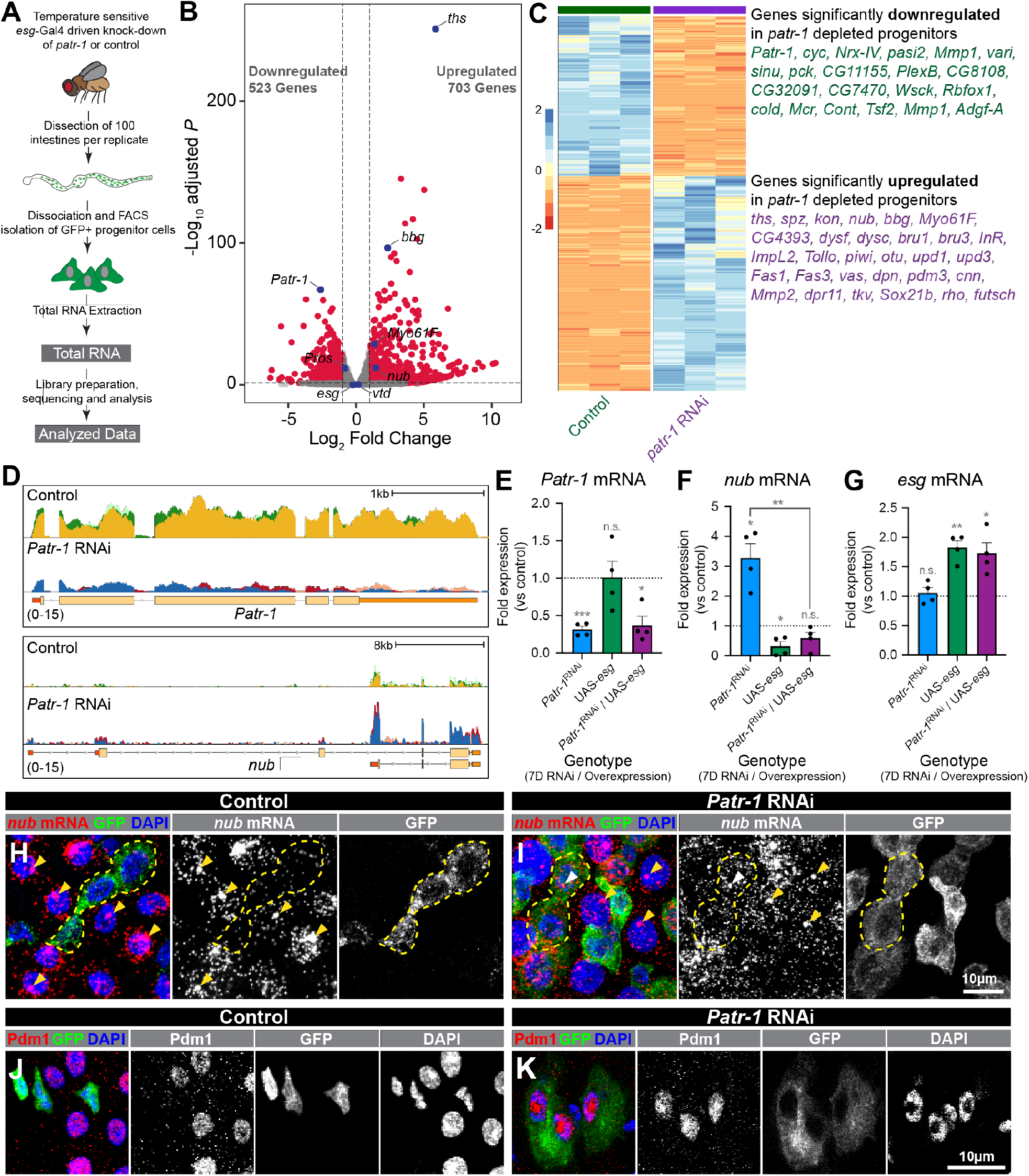
PATR-1 limits pro-differentiation gene expression in intestinal progenitor cells. (A) Schematic of RNA-seq analysis. (B) Volcano plot of differentially expressed genes in *Patr-1* RNAi versus control. Each dot represents a single gene. Red indicate a false discovery rate (FDR) adjusted p-value < 0.05 and a Log_2_ fold change > 1 or < −1. (C) Heatmap showing differentially expressed genes (FDR < 0.05; Log_2_ fold change > 1 or < −1) in *Patr-1* RNAi versus control with a selected set of gene names shown. (D) Genome browser tracks of CPM-normalized reads from control and *Patr-1* RNAi at the *Patr-1* and *nub* loci. Replicates of control (yellow, light green and dark green) and *Patr-1* RNAi (blue, pink and red) have been overlayed. (E-G) Fold expression of (E) *Patr-1*, (F) *nub*, and (G) *esg* mRNA in intestinal progenitors of *esg*^*TS*^ */ Patr-1 RNAi, esg*^*TS*^ / *UAS-esg* and *esg*^*TS*^ */ Patr-1 RNAi* / *UAS-esg* compared to *esg*^*TS*^ as determined by qPCR. Error bars show mean±s.d. and asterisks denote statistical significance from one sample t and Wilcoxon test \**p* < 0.05; \*\**p* < 0.01; n.s., not significant. (H-K) Intestinal progenitors from (H, J) *esg*^*TS*^ and (I, K) *esg*^*TS*^ */ Patr-1 RNAi* stained for (H-I) *nub* mRNA (red) or (J-K) Pdm1 (red) and GFP (green) and DAPI (blue). Yellow dotted outline on H-I marks intestinal progenitor cells. Yellow and white arrowheads on H-I mark nuclear staining that we interpret as active transcription in ECs and progenitors, respectively.

### PATR-1 limits cytoplasmic *nub* mRNA levels in progenitor cells

To evaluate the relationship between transcriptional and post-transcriptional regulation in progenitor cells, we focused on the *nub* mRNA, which we confirmed via real-time quantitative PCR (qPCR) was upregulated in FACS-isolated *Patr-1* mutant cells (**Fig. 4E-F**). This qPCR analysis, however, detected low levels of *nub* in wildtype intestinal progenitors, suggesting that repression of these transcripts in wildtype progenitors was incomplete. This qPCR result was consistent with low levels of *nub* and other pro-differentiation genes including *bbg* detected by RNA-seq analysis of wildtype progenitors (**Fig. 4D**). To directly confirm these observations, we analyzed the subcellular distribution and levels of *nub* transcript in wildtype intestines using RNAscope *in situ hybridization* probes. As expected, *nub* transcript was readily detected in ECs in both the cytoplasm as well as in bright nuclear punctae that we interpreted as sites of active transcription (**Fig. 4H**, yellow arrowheads). In addition, *nub* transcript was more weakly detected in the cytoplasm of ~33% wildtype progenitor cells, with low-to-no signal in the nuclei of these cells (**Fig. 4H**). In comparison, the *nub* protein product Pdm1, was rarely detected in only ~3% of wildtype progenitors (**Fig. 4J**, quantified in **5A)**. These results indicated that the transcriptional repression of *nub* was incomplete and that a post-transcriptional mechanism ensured the absence of Pdm1 protein in wildtype progenitor cells.

To analyze the role of P-bodies in this process, we also analyzed *nub* mRNA and Pdm1 protein patterns and levels in intestinal progenitor cells after *Patr-1* knockdown. In comparison to wildtype progenitors, cytoplasmic *nub* mRNA staining was considerably higher in ~63% *Patr-1* deficient intestinal progenitors (**Fig. 4I**). Only 40% subset of the *Patr-1* mutant progenitor cells with elevated cytoplasmic staining also displayed bright nuclear punctae (**Fig. 4I**, white arrowhead), indicating that elevated cytoplasmic expression did not require nuclear expression. Furthermore, neither *esg* nor *vtd* expression levels were affected by *Patr-1* loss (**Fig. S4H-I**), suggesting that transcriptional repression was unlikely to be compromised by P-body loss. In addition, the number of progenitor cells expressing Pdm1 protein was significantly enhanced after *Patr-1* RNAi (**Fig. 4K**, quantified in **5A**). Collectively, these observations indicated that PATR-1 mediated P-body assembly in progenitor cells was required to limit the stability and translation of pro-differentiation genes such as *nub*.

### Loss of P-body assembly promotes progenitor-EC differentiation

The elevated Pdm1 protein and transcript levels detected in *Patr-1* mutant progenitor cells suggested that P-body loss caused premature progenitor-to-EC differentiation. Consistently, forced expression of Pdm1 in progenitor cells also led to progenitor cell loss (**Fig. 5B)**. In addition, prior to their loss, both the cell and nuclear area of *Patr-1* mutant intestinal progenitors was significantly elevated (**Fig. 5C-D**), suggestive of the acquisition of polyploid EC characteristics. Since no increase in GFP+ PROS+ cells or apoptotic cell death was observed (**Fig. S5A-C)**, these observations indicated that P-body loss was associated with derepressed translation of pro-differentiation gene products in progenitor cells that led to progenitor-to-EC differentiation and consequent progenitor cell loss. Consistently, we noted that feeding adults bleomycin, which leads ISCs to rapidly proliferate, differentiate and upregulate Pdm1 (Tian et al., 2017), was also associated with P-body dissolution (**Fig. S5D-I**). Taken together, these data showed that P-bodies are required for progenitor cell maintenance by suppressing EC-fate.

**Figure 5:**
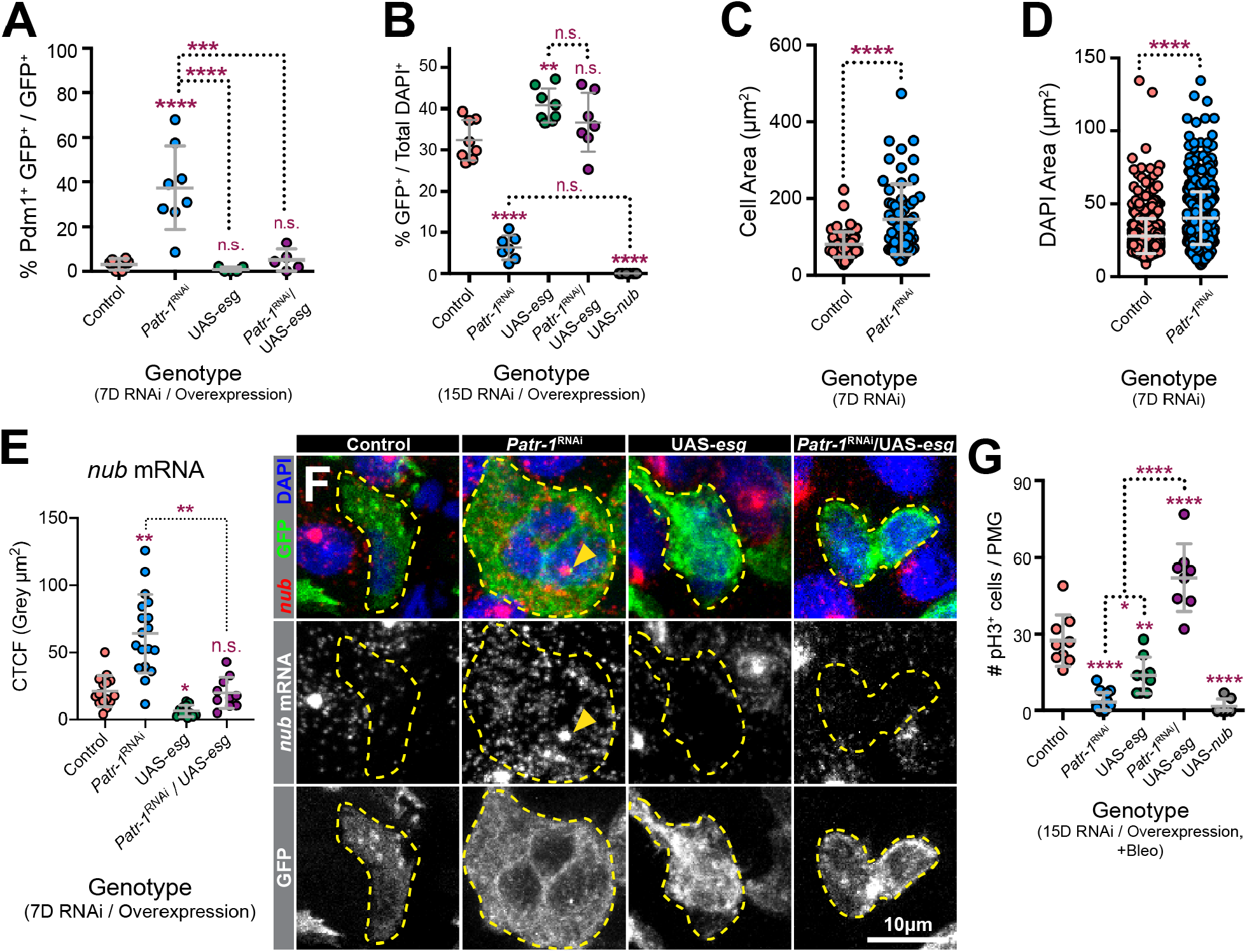
Loss of PATR-1 promotes progenitor-to-EC-differentiation. (A-E) Dot-plots showing (A) percentage of intestinal progenitors with nuclear Pdm1 staining (n=6, 8, 5, or 5), (B) normalized intestinal progenitor cell percentage (n=8, 7, 8, 7, or 8), (C) cell area (n=58 or 61), or (D) nuclei area (n=717 or 619) of indicated genotypes after (A, C, D) 7 or (B) 15 days at 29°C. (E) Dot plot showing the normalized *nub* mRNA fluorescence intensity of progenitor cells from *esg*^*TS*^ (n=15 cells), *esg*^*TS*^ */ Patr-1 RNAi* (n=18 cells), *esg*^*TS*^ */ UAS-esg* (n=12 cells), or *esg*^*TS*^ */ Patr-1 RNAi* / *UAS-esg* (n=10 cells) intestines. (F) Intestinal progenitors from genotypes in E stained for *nub* mRNA (red), GFP (green) and DAPI (blue). Yellow dotted outline marks intestinal progenitor cells. Yellow arrowhead mark nuclear staining that we interpret as active transcription. (G) Bleomycin-induced pH3+ cell number per posterior midgut (n=9, 14, 9, 8, or 7) of indicated genotypes after 15 days at 29°C. Error bars on plots show mean±s.d. and asterisks denote statistical significance from Ordinary one-way ANOVA with Turkey’s multiple comparison test (A-B, G), Mann-Whitney test (C-D) or Kruskal-Wallis test (E). \**p* < 0.05; \*\**p* < 0.01; \*\*\*\**p* < 0.0001; n.s., not significant.

### Overexpression of *esg* rescues *Patr-1* RNAi mediated progenitor cell loss

The presence of *nub* transcript but absence of Pdm1 protein in progenitor cells suggested that P-bodies limited the translation of weakly transcribed pro-differentiation genes to maintain progenitor cell identity. To test whether the overexpression of *esg*, which we hypothesized would block this weak transcription, eliminated the requirement of mature P-body formation to prevent differentiation, we analyzed the effects of progenitor expression of UAS-*Patr-1* RNAi and UAS-*esg* transgenes, both individually and in combination, on progenitor cell numbers. Consistent with our previous observations, knocking down *Patr-1* significantly reduced the number of intestinal progenitors while *esg* overexpression significantly increased the progenitor cell number (**Fig. 5B, S5J**). However, the ectopic expression of Pdm1 and ensuing progenitor cell loss associated with *Patr-1* knockdown was eliminated when *esg* was simultaneously expressed (**Fig. 5A-B, S5J-K**). We verified this at the mRNA level with both qPCR and RNAscope analysis. As expected, *esg* but not *Patr-1* transcript abundance was significantly elevated in progenitors expressing UAS-*esg* (**Fig. 4E, G**). Consistent with our model, this ectopic *esg* expression significantly reduced *nub* mRNA level in both wildtype and *Patr-1* mutant progenitor cells (**Fig. 4F, 5E-F**). To verify that the rescued progenitor cells in the UAS-*Patr-1* RNAi/UAS-*esg* background included bona fide ISCs, we assessed the mitotic potential of intestines of each of the above genetic backgrounds following bleomycin feeding. As expected, overexpression of *esg* in *Patr-1* deficient intestinal progenitors elevated the total number of mitotic ISCs (**Fig. 5G**). However, mature P-bodies were not restored, as indicated by diffuse TRAL staining in these intestinal progenitors (**Fig. S5L**). Taken together, enhanced transcriptional repression of pro-differentiation genes associated with *esg* overexpression can prevent premature progenitor differentiation independent of mature P-body assembly.

## Discussion

This study molecularly and functionally characterized stem cell mRNPs that we conclude are P-bodies based on three observations: (i) they contain colocalized protein complexes that include fly orthologs of proteins known to localize to P-bodies in mammalian and yeast cells, (ii) these mRNP granules are significantly larger than and show no colocalization with IPSG protein foci under controlled conditions, and (iii) acute stresses increase the size of these mRNPs and promotes colocalization with IPSGs. A targeted genetic screen identified 103 genes, 39 previously known and 64 novel candidates, that can influence P-body morphology, including six that are required for P-body formation. To examine stem cell P-body function, we genetically characterized one of these latter class, PATR-1, an evolutionary conserved protein with both translational repression and mRNA decay functions that is necessary for proper P-body assembly in *Saccharomyces cerevisiae* (Coller and Parker, 2005; Eulalio et al., 2007b; Scheller et al., 2007; Sheth and Parker, 2003; Teixeira and Parker, 2007) (Pilkington and Parker, 2008; Sachdev et al., 2019). Genetic depletion of P-bodies in progenitor cells upregulated the expression of pro-differentiation genes such as *nubbin*, which was weakly transcribed but translationally repressed in wildtype progenitor cells. Loss of stem cell P-bodies, either by genetic depletion or differentiation, led to the increased translation as well as the cytoplasmic, but not nuclear, abundance of such transcripts.

We therefore concluded that mature P-bodies are necessary for stem cell maintenance by post-transcriptionally enforcing the repression of transcriptional programs that promote differentiation. Differentiation signals dissolve P-bodies and thereby promote translation of target differentiation genes (**Fig. 6**).

**Figure 6:**
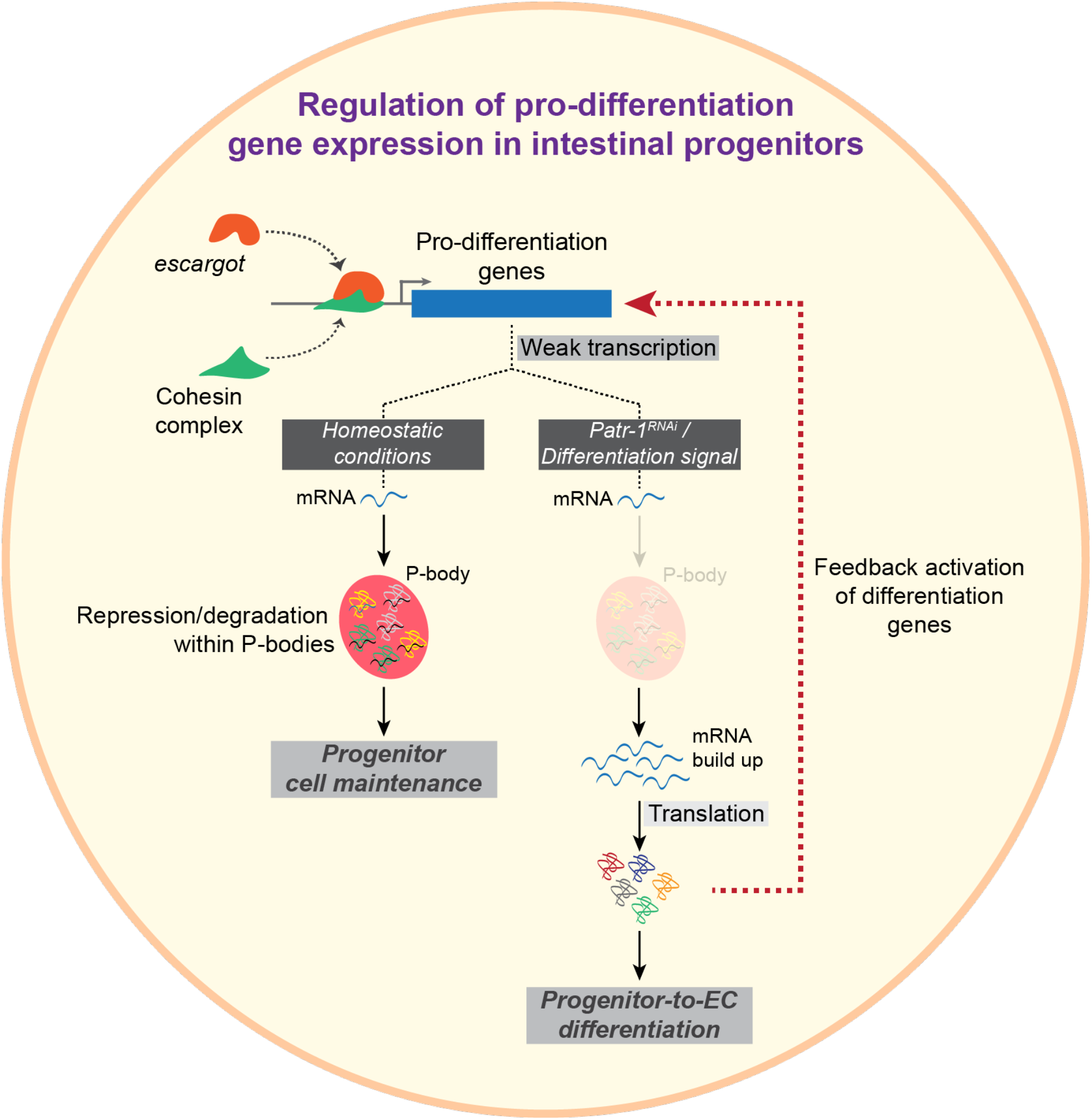
P-body mediated post-transcriptional regulation maintains progenitor cell identity. The model proposing how Patr-1 mediated P-body formation modulate progenitor cell fate.

Since detailed ultrastructural organization of P-bodies has been lacking (Youn et al., 2019), we used quantitative super-resolution microscopy-based characterization to show substructures present within mature P-bodies. Consistent with the proposed “core-shell” structure of stress granules (Jain et al., 2016; Niewidok et al., 2018; Wheeler et al., 2016; Youn et al., 2019), P-bodies exhibit “cores” with high protein concentrations and “shells” with low protein concentration. Notably, Drosophila intestinal progenitor P-bodies have a diameter of ~125 nm, as compared to the larger P-bodies in HEK293 cells, which have a ~500 nm diameter (Hubstenberger et al., 2017). In addition, intestinal progenitors contain ~35-45 mature P-bodies while HEK293 cells contain only ~4-7 granules per cell. This indicates that the size and the number of mature P-bodies depends on cell type and species and may scale based on overall cell size.

A recent study documented the presence of P-bodies in cultured human pluripotent stem cells (hPSCs) and suggested their presence in adult stem cells (Di Stefano et al., 2019). This analysis of DDX6, the ortholog of *Drosophila* Me31B, showed that DDX6-dependent P-bodies could both promote and repress stem cell identity, depending on context. For example, loss of DDX6 expanded endodermally derived Lgr5^+^ ISC or ectodermally derived neural progenitor cell populations, but promoted the differentiation of other progenitor cell populations, including mesodermally derived progenitors. We confirm the presence of mature P-bodies in adult progenitor populations but show that they repress differentiation rather than increasing their proliferation, as in Lgr5^+^ ISCs. A few possible explanations could explain these results. Most simply, *Drosophila* intestinal progenitors behave more like mesodermally derived mammalian progenitors rather than endo- or ectodermally derived mammalian progenitors. Alternatively, the stem cell function of DDX6 might be affected by its roles in surrounding cells, since DDX6 was targeted in cells throughout mouse intestinal organoids, whereas PATR-1 was specifically targeted only in progenitor cells in our study. Finally, DDX6-mediated P-body function might be modulated by signaling that is not fully recapitulated in *in vitro*-derived stem cell models.

The exact molecular function of P-bodies is a matter of current debate (Sachdev et al., 2019). While earlier models proposed that the primary function of P-bodies was mRNA degradation, more recent findings propose that P-bodies function as sites of mRNA storage and control the translatome of cells (Anderson and Kedersha, 2006; Decker and Parker, 2012; Standart and Weil, 2018). While our RNA-seq analysis identifies potential targets of P-bodies, more direct approaches such as enhanced-CLIP (eCLIP) (Van Nostrand et al., 2016) of P-body proteins driven with recently developed progenitor specific *I-KCKT-GAL4* drivers (Buddika et al., 2020b) will be used in the future to directly profile the P-body transcriptome of intestinal progenitors. Our data, however, suggest that intestinal progenitor P-bodies may have both translational repressive and mRNA degratory functions. We found that the *nub* mRNA was weakly expressed but not translated in wildtype intestinal progenitors, and that this translational repression was P-body dependent. In addition, the abundance of EC-specific transcripts was influenced by P-bodies in a transcription independent fashion: RNAscope *in situ* analysis showed that the cytoplasmic *nub* transcript abundance was increased in *Patr-1* mutant progenitor cells without an indication of nuclear transcription. Therefore, this PATR-1 dependent increase in cytoplasmic *nub* mRNA could be attributed to the stabilization of transcripts that are targeted for degradation via P-bodies. Consistently, studies of P-bodies and P-body proteins in human cultured stem cells also sugget dual roles in both mRNA translation repression and degradation. For instance, P-bodies influence the translation of transcripts encoding fate-instructive transcription and chromatin factors in cultured embryonic and *in vitro*-derived adult stem cells (Di Stefano et al., 2019). In addition, P-body proteins DDX6 and EDC3 are known to destabilize differentiation-inducing mRNAs such as KLF4 in human epidermal progenitor cells, although it is important to note that P-bodies have not been reported in these cells (Wang et al., 2015).

Based on our data, the weak transcription of pro-differentiation genes likely maintains progenitor cells in a primed state where differentiation program can be initiated rapidly and independent of transcription of differentiation genes. The transcriptional repression of differentiation genes by the transcription factor *esg* is a key regulatory step of intestinal stem cell maintenance (Antonello et al., 2015; Khaminets et al., 2020; Korzelius et al., 2014; Li et al., 2017; Liu and Jin, 2017; Loza-Coll et al., 2014). The loss of *esg* gene expression or inability to localize the Esg protein to target genes promote progenitor cell loss via premature ISC-to-EC differentiation (Khaminets et al., 2020; Korzelius et al., 2014). Similar to *Patr-1* RNAi, knocking down *esg* itself as well as either *vtd, Nipped-B* or *polo*, all of which are necessary for recruiting Esg to target promoters, markedly upregulated the expression of Pdm1 in intestinal progenitors (Khaminets et al., 2020; Korzelius et al., 2014). Notably, our transcriptomic profiling showed that the transcript level of neither *esg* nor any of the Esg-targeting proteins, *vtd, Nipped-B* or *polo*, was changed by the absence of mature P-bodies. These observations suggest that Esg protein level, its proper promoter targeting, and its transcriptional repression of EC-genes are all unlikely to be affected by the loss of PATR-1.

In addition to identifying 64 new genes affecting P-body morphology, we expect that the tissue-based stem cell P-body system identified and described here will prove critically useful in screening for chemicals, diet conditions and stress conditions that alter P-body assembly as well as performing larger, genome-wide screens to comprehensively characterize the molecular pathways that control P-body assembly. Moreover, similar approaches can be used to identify systemic signals that promotes P-body disassembly during the onset of differentiation as well as to identify molecular players of P-body disassembly.

## Acknowledgements

We thank Scott Barbee, Anne Ephrussi, Xiaohang Yang, the Bloomington *Drosophila* Stock Center (supported by grant NIHP4OOD018537), the *Drosophila* Genome Resource Center (supported by grant NIH2P40OD010949), and the Developmental Studies Hybridoma Bank (created by the NICHD of the NIH) for reagents; Robert Policastro for bioinformatics expertise; the Light Microscopy Imaging Center (supported by grant NIH1S10OD024988-01) for access to the SP8 confocal and OMX super-resolution microscopes; the Flow Cytometry Core Facility at Indiana University, Bloomington for access to the BD FACSAria™ II flow cytometer; and the National Institute of General Medical Sciences (Award R01GM124220) for financial support.

## STAR Methods

Key reagents and resources are listed in **Table S3**.

### Drosophila strains and husbandry

Age matched female flies were used in all experiments. Fly strains were cultured on standard Bloomington *Drosophila* stock center media (https://bdsc.indiana.edu/information/recipes/bloomfood.html) and reared in 18°C, 25°C and 29°C incubators set for a 12hr light/dark schedule and 65% humidity. Flies were cultured in groups of 15-20 (typically 5 males and up to 15 females). All strains used in individual panels are listed in **Table S4**.

### Temperature

For temporal and regional gene expression-targeting (TARGET) experiments, flies were grown at 18°C, collected over 2 days, and reared in 29°C for up to 20 days before being dissected. For clonal analysis using mosaic analysis with repressible cell marker (MARCM) method, animals were reared at 25°C until eclosion, collected over 2 days and heat-shocked immediately at 37°C for 45 min in a Lauda circulating water bath.

Subsequently, flies were reared at 25°C for 7-15 days.

### Construction of new strains

#### CRISPR/cas9 mediated generation of Patr-1 alleles

New *Patr-1* alleles were generated by co-injection of two guide RNA (gRNA) plasmids. The gRNA plasmids were generated by subcloning annealed oligos encoding gRNA 5’-catgttgaacatgttatacatgg-3’ as well as annealed encoding gRNA 5’-tgtgacgagactgtcggaagggg-3’ into the BbsI site of *pU6-BbsI-chiRNA* (Gratz et al 2013). The two resulting plasmids were sequence-verified, amplified, mixed, and co-injected into strain *y*^*1*^ *sc* v*^*1*^ *sev*^*21*^; *P{nos-Cas9}attP2* (BL78782) by Rainbow Genetics (Camarillo CA). Stocks of F_1_ progeny were generated, and strains containing new *Patr-1* alleles were selected based on non-complementation of the lethality associated with *P{EPgy2}Patr-1*^*EY10289*^ (BL19805). New *Patr-1* alleles were molecularly defined by sequencing PCR products generated from genomic DNA obtained from homozygous mutant larvae with oligos flanking the gRNA locations. Two new *Patr-1* alleles as well as *P{EPgy2}Patr-1*^*EY10289*^ were recombined onto P{FRT}^82B^-containing chromosomes using standard meiotic methods.

### Antibody generation

Anti-TRAL antibodies were generated in rats (Cocalico Biologicals, Reamstown PA) against a 6XHIS-tagged version of an N-terminal portion of TRAL that was expressed and purified according to standard methods. The TRAL-encoding plasmid was generated by PCR amplifying a 477bp fragment that encodes the first 160 amino acids of TRAL from cDNA GH08269 (DGRC stock 4838) with high-fidelity Q5 polymerase (NEB), subcloning the resulting PCR product into the NcoI and EcoRI sites of pHIS.parallel using HiFi DNA Assembly Master Mix (NEB), and sequence-verifying the resulting plasmid to confirm the absence of any PCR-induced errors.

Anti-EDC3 antibodies were generated in rabbits and rats (MBL, Japan) against a 6XHIS-tagged version of an N-terminal portion of EDC3. The *edc3* coding sequence (corresponding to amino acid 1-440) was PCR-amplified from an ovarian cDNA library using Phusion DNA polymerase (Finnzymes), with primers 5’-GGGAATTCATGGGTCCGACGGATCAAGA-3’ and 5’-GGGGCGGCCGCTCACTTATCGGCACTTATCTCGA-3’. The fragment was digested by *Eco*RI and *Not*I, and cloned into the pProEX HTa vector (Life Technologies). The 6×His-tagged Edc3 (1-440) protein was expressed in *E. coli* BL21 cells by IPTG induction, and purified using Ni-NTA agarose resin (Qiagen) under denaturing conditions. The protein was further purified by disc preparative SDS-PAGE using the NA-1800 apparatus (Nihon Eido, Japan). Purified protein fractions were concentrated with Vivaspan-2 (10,000 MWCO PES; Sartorius), dialyzed against PBS containing 4M urea followed by PBS/2M urea. Polyclonal antibodies were affinity-purified with the same antigen immobilized on HiTrap NHS columns (GE Healthcare).

### Dissections and immunostaining

Gastrointestinal (GI) tracts of adult female flies were dissected in ice-cold 1xPBS and fixed in 4% paraformaldehyde (Electron Microscopy Sciences, Cat. No. 15714) in PBS for 45 min. These samples were washed with 1xPBT (1xPBS, 0.1% Triton X-100) and then blocked (1xPBT, 0.5% Bovine Serum Albumin) for at least 45 min. Subsequently, samples were incubated at 4°C overnight with primary antibodies, including rabbit anti-GFP (A11122, Life Technologies, 1:1000), mouse anti-V5 (MCA1360GA, Bio-Rad, 1:250), mouse anti-FLAG (F3165, Sigma, 1:1000), rabbit anti-HA (3724S, Cell Signaling Technology, 1:1000), mouse anti-FMR1 (5A11, Developmental Studies Hybridoma Bank, 1:100), mouse anti-Prospero (MR1A, Developmental Studies Hybridoma Bank, 1:100), mouse anti-Delta (C594.9B, Developmental Studies Hybridoma Bank, 1:500), rat anti-TRAL (this study, 1:1500), rabbit anti-Me31B (Nakamura A, Amikura R, Hanyu K, 2001) (1:2000), mouse anti-Me31B (Barbee et al., 2006) (1:1500), rabbit anti-Ge-1 (Fan et al., 2011) (1:500), rat anti-EDC3 (this study, 1:1500), rabbit anti-EDC3 (this study, 1:2000) and rabbit anti-Pdm1 (a gift from Xiaohang Yang) (1:1500). Samples were washed and incubated for 2-3 hours with secondary antibodies, including AlexaFluor-488 and -568-conjugated goat anti-rabbit, -mouse and -rat antibodies (Life Technologies, 1:1000). AlexaFluor-647 conjugated goat-HRP antibodies were used in the secondary antibody solution whenever required. Samples were washed and treated with DAPI (1:10000) and mounted in Vectashield (Vecta Laboratories). An alternative staining protocol was used for Delta staining as described in Buddika et al., 2020a.

### RNAscope in situ hybridization

Adult female flies were dissected in ice cold 1X PBS and fixed in 4% paraformaldehyde (Electron Microscopy Sciences, Cat. No. 15714) in PBS for 45 min. Tissue was then washed in 1xPBT (1xPBS, 0.3% Triton X-100) 3 times 5 min each. Next, samples were gradually dehydrated with a series of 0.3% PBT (1xPBS, 0.3% Triton X-100): Methanol (7:3, 1:1, 3:7) washes and incubated in Methanol for 10min. Then tissue was rehydrated with a series of 0.3% PBT: Methanol (3:7, 1:1, 7:3) washes and washed with 0.3% PBT alone for another 5 min. For following steps, reagents from RNAscope Multiplex Fluorescent Reagent Kit v2 assay were used. Fixed tissue was transferred into 0.2 ml PCR tubes and incubated in RNAscope Protease III reagent for 5 min at 40°C (PCR thermal cycler was used for all the incubations at 40°C). Samples were immediately washed with 1X PBS twice and 20µl of RNAscope probes for *nub* (pre warmed to 40°C) (ACD Bio., Cat. No. 523981) was added. Samples were incubated at 40°C overnight. After overnight incubation, samples were washed twice with 1X RNAscope wash buffer. Next, RNAscope Multiplex FL v2 AMP 1, RNAscope Multiplex FL v2 AMP 2, RNAscope Multiplex FL v2 AMP 3 and RNAscope Multiplex FL v2 HRP-C1 steps were done as described in Chapter 4 of Fluorescent v2 assay manual, ACD Bio. Finally, samples were incubated for 30 min at 40°C with Opal 620 (AKOYA Biosciences, Cat. No. FP1495001KT, 1:1500 in TSA buffer) and washed with 1X RNAscope wash buffer and counterstained with DAPI. Samples were mounted in ProLong Diamond mounting medium (Invitrogen).

### Microscopy and image processing

Images were collected on either a Leica SP8 Scanning Confocal microscope (Leica DMi8 inverted microscope platform; equipped with WLL 470-670 nm, 405 nm and 440 nm lasers, Huygens deconvolution software; controlled by Leica LAS-X software; image acquisition using Leica HC PL APO CS2 63x/1.40 or Leica HC PL APO CS2 40x/1.40 lens with Leica Type F Immersion Liquid (N = 1.518)) or an OMX 3D-SIM Super-Resolution microscope (DeltaVision OMX system; equipped with 405, 488, 561, 642 nm lasers; controlled by AquireSR software; image processing by SoftWorx imaging software; image acquisition using an Olympus PL APO N 60x/1.42 lens with Applied Precision Immersion Oil (N = 1.518)) available at the Light Microscopy Imaging Center, Indiana University, Bloomington. Whenever possible, samples to be compared were collected under identical settings on the same day, image files were adjusted simultaneously using Adobe Photoshop CC, and figures were assembled using Adobe Illustrator CC. ImageJ FIJI (https://fiji.sc/) Analyze Particles plugin was used to quantify the average area of granules using 0.01µm^2^ as the lower cut off of puncta size. The ImageJ FIJI Plot Profile plugin was used to generate line plots.

### Ex vivo treatments

All *ex-vivo* treatments were performed as described in Buddika et al., JCS, 2020. Briefly, intestines from females aged 8-10 days on normal diet were incubated in Krebs-Ringer media/KRB (Alfa Aesar, Cat. No. J67591) or KRB supplemented with 1mM rapamycin (LC Laboratories, Cat. No. r-5000) for 60 min and then fixed.

### Bleomycin feeding assay

Female flies were maintained on standard Bloomington *Drosophila* stock center media for an appropriate time and then separated into two cohorts. Cohort 1 and 2 were transferred to a vial with a chromatography paper soaked in either 5% sucrose in water (control) or 5% sucrose and 25 µg/ml bleomycin in water, respectively. Based on experiments, flies were dissected 24-to-36 hours after feeding.

### Protein isolation and western blot analysis

Wandering L3 larvae or adult female flies were used for protein isolation. Collected larvae or adult flies were lysed in I-RIPA protein lysis buffer (150mM NaCl, 50mM Tris-HCl pH 7.5, 1mM EDTA, 1% Triton X-100, 1% Na Deoxycholic Acid, 1xprotease inhibitor cocktail), protein extracts were resolved on a 4-20% gradient polyacrylamide gel (Bio-Rad, Cat. No. 456-1093), transferred to Immobilon^®^-P membrane (Millipore, Cat. No. IPVH00010) and probed with rat anti-TRAL (1:2000, this study), rat anti-PATR1 (Pradhan et al., 2012) (1:2000) or mouse anti-a-tubulin (12G10, Developmental Studies Hybridoma Bank, 1:1000) antibodies. Subsequently, blots were washed extensively with 1xTBST (1xTBS, 0.1% Tween-20) and incubated with anti-rat or -mouse conjugated HRP secondary antibodies. After extensive secondary washes with 1xTBST, blots were treated with ECL-detection reagent 1 and 2 (Thermo Scientific, Cat. No. 1859701 and 1859698) and finally exposed to chemiluminescence films (GE Healthcare, Cat. No. 28906839) and developed the signal.

### FACS isolation of progenitor cells, RNA-seq library preparation and qPCR

Gastrointestinal (GI) tracts of 100 adult female flies (per replicate) were dissected in ice-cold 1xPBS. Then cells were dissociated by treating intestines with 1mg/ml elastase at 27°C for 1 hour with agitation. Subsequently, ~30,000-50,000 GFP^+^ intestinal progenitor cells were sorted using a BD FACSAria™ II flow cytometer equipped with a 100µm nozzle at the IUB Flow Cytometry Core Facility. Total RNA was prepared using the TRIzol® LS reagent (Ambion). The rRNA-depleted libraries were prepared using the Ovation® SoLo RNA-seq system (Part No. 0502 including Parts 0407 and S02240) following manufacturer’s instructions. The quality and quantity of final libraries were assessed using Agilent 2200 TapeStation and KAPA Library Quantification Kit, respectively. For qPCR, resulting RNA was first treated with Turbo DNase (ThermoFisher, AM2239) and gDNA-free RNA used for cDNA synthesis with Superscript III (ThermoFisher, 56575). qPCR was performed using the PowerUp SYBR Green Master Mix (ThermoFisher, A25742) on a StepOnePlus machine (ThermoFisher). Primers for all targets detected are listed in **Table S3**. Transcript levels were quantified in triplicates and normalized to *Gapdh1*. Fold enrichment was calculated as the ratio of transcript in genetic manipulation versus control.

### RNA-seq data analysis

Transcriptomic data analysis was performed as described in Buddika et al., 2020a using a python based in-house pipeline (https://github.com/jkkbuddika/RNA-Seq-Data-Analyzer). Briefly, the quality of raw sequencing files was assessed using FastQC (Andrews, 2010) version 0.11.9, low quality reads were eliminated using Cutadapt (Martin, 2011) version 2.9, and reads mapping to rRNAs were removed using TagDust2 (Lassmann, 2015) version 2.2. Next, the remaining reads were mapped to the Berkeley *Drosophila* Genome Project (BDGP) assembly release 6.28 (Ensembl release 100) reference genome using STAR genome aligner (Dobin et al., 2013) version 2.7.3a and duplicated reads were removed using SAMtools (Li et al., 2009) version 1.10. Subsequently, the Subread (Liao et al., 2019) version 2.0.0 function *featureCounts* was used to count the number of aligned reads to the nearest overlapping feature. Finally, bigWig files representing RNA-seq coverage were generated using deepTools (Ramírez et al., 2016) version 3.4.2 with the settings --normalizeUsing CPM --binSize 1. Differential gene expression analysis was performed with the Bioconductor package DESeq2 (https://bioconductor.org/packages/release/bioc/html/DESeq2.html) (Love et al., 2014) version 1.26.0. Unless otherwise noted, significantly upregulated and downregulated genes were defined as FDR < 0.05; Log_2_ fold change > 1 and FDR < 0.05; Log_2_ fold change < −1, respectively and were used to identify enriched Gene Ontology (GO) terms using PANTHER overrepresentation analysis on GO Consortium (http://geneontology.org/). A selected significantly enriched GO categories were plotted. All data visualization steps were performed using custom scripts written using R.

### Statistical analysis

GraphPad Prism, Version 7.0 was used for all statistical analyses. First, the normality of datasets was tested using D’Agostino-Pearson test. For comparisons involved in two datasets, if datasets follow [1] a parametric distribution, an Unpaired t-test or [2] a non-parametric distribution, a Mann-Whitney test was performed. Three or more datasets following a parametric distribution were analyzed using an ordinary one-way ANOVA test. Multiple comparisons of three or more datasets following a non-parametric distribution were analyzed using Kruskal-Wallis test. Unless otherwise noted, significance is indicated as follows: n.s., not significant; *p < 0.05; **p < 0.01; ***p < 0.001; ****p < 0.0001.

### Data availability

The control and *Patr-1* RNAi RNA-seq datasets from this study have been submitted to the NCBI Gene Expression Omnibus under accession number XXXXXX.

## Supplemental Information and Legends

**Figure S1:**
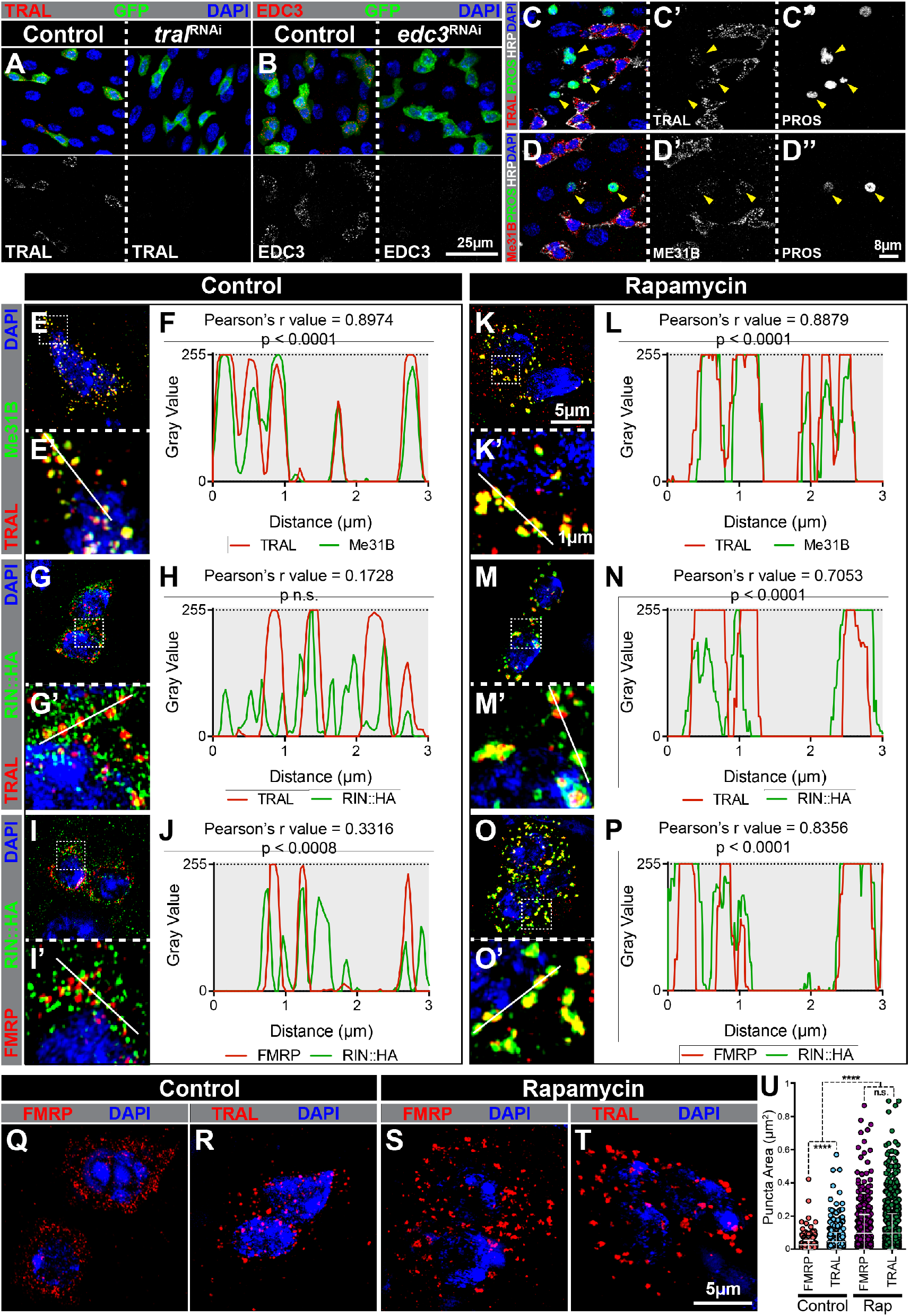
Comparative analysis of P-body and stress granule component expression. (A-B) Progenitor cells from (A) *esg*^*TS*^ (control, left), *esg*^*TS*^ */ tral* RNAi (right) or (B) *esg*^*TS*^ (control, left), *esg*^*TS*^ */ edc3* RNAi (right) intestines stained for (A) TRAL (red) or (B) EDC3 (red). Note that *tral* or *edc3* mutant GFP+ intestinal progenitors have no detectable TRAL or EDC3 staining, respectively. (C-D) Confocal micrographs of intestinal sections stained for (C) TRAL (red) or (D) Me31B (red), (C-D) PROS (green), (C-D) HRP (white) and (C-D) DAPI. While TRAL and Me31B granules are readily observed in all HRP+ intestinal progenitors, only some Prospero+ EEs have detectable TRAL or Me31B expression. (E, G, I, K, M, O) Super-resolution micrographs of progenitor cells treated with either (E, G, I) KRB or (K, M, O) 1mM rapamycin in KRB and stained for (E, K) TRAL (red), Me31B (green) or (G, M) TRAL (red), RIN::HA (green) or (I, O) FMRP (red), RIN::HA (green) and DAPI (blue in E, G, I, K, M, O). E’, G’, I’, K’, M’, and O’ show a magnified view of the region marked with white dotted outline in E, G, I, K, M, and O respectively. (F, H, J, L, N, P) Line plots of fluorescent intensity profiles of (F, L) TRAL (red), Me31B (green) or (H, N) TRAL (red), RIN::HA (green) or (J, P) FMRP (red), RIN::HA (green). The plot shows the pixel-by-pixel gray value of each channel along white lines in E’, G’, I’, K’, M’ or O’. Computed Pearson’s r value and the statistical significance of the correlation are given above each plot. Note that Me31B and TRAL are P-body proteins while FMRP and RIN::HA are stress granule proteins. (Q-T) Super-resolution micrographs of progenitor cells treated with either (Q-R) KRB (control) or (S-T) 1mM rapamycin in KRB, and stained for (Q, S) FMRP (red) or (R, T) TRAL (red) and (Q-R) DAPI (blue). (U) Scatter dot plots of FMRP or TRAL puncta area in progenitor cells treated with control (n=629 or 657) or 1mM rapamycin (n=804 or 1215) in KRB. Error bars on plots show mean±s.d. and asterisks denote statistical significance from Kruskal-Wallis test (I) \*\**p* < 0.01; \*\*\*\**p* < 0.0001; n.s., not significant.

**Figure S2:**
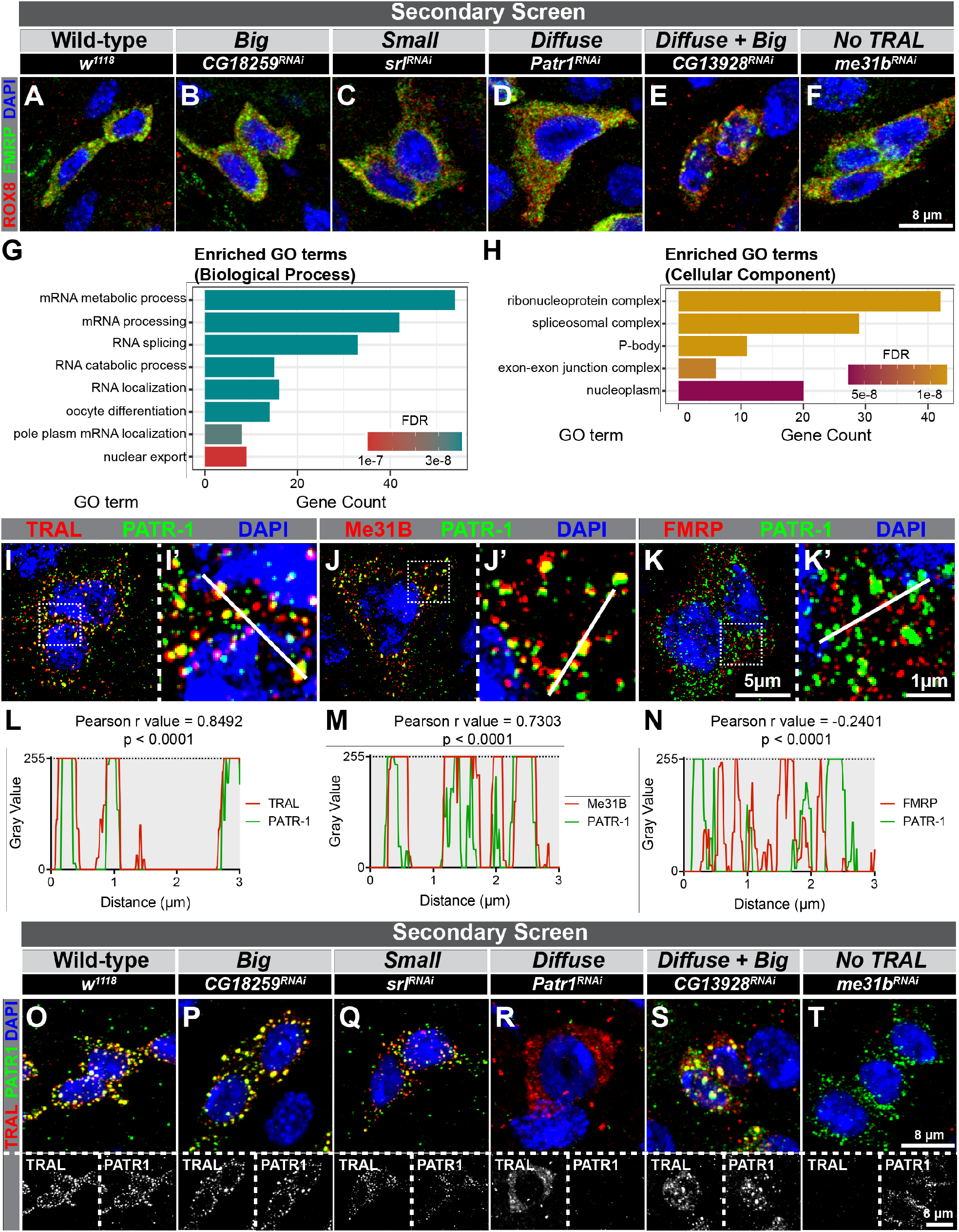
A targeted genetic screen identifies genes affecting P-body morphology including the P-body component, PATR-1. (A-F) Confocal micrographs of intestinal progenitors from (A) *esg*^*TS*^, (B) *esg*^*TS*^ / *CG18259* RNAi, (C) *esg*^*TS*^ / *srl* RNAi, (D) *esg*^*TS*^ / *Patr-1* RNAi, (E) *esg*^*TS*^ / *CG13928* RNAi or (F) *esg*^*TS*^ / *me31b* RNAi posterior midguts and stained for ROX8 (red), FMRP (green) and DAPI (blue). (G-H) Bar plots visualizing significantly enriched (G) biological process or (H) cellular component Gene Ontology (GO) terms (FDR < 0.05) for positive genes. The y-axis represents significantly enriched GO term and the x-axis represent gene count for each GO term. Color of each bar represent the FDR. (I-K) Super-resolution micrographs of intestinal progenitors stained for (I-I’) TRAL (red), PATR-1 (green) or (J-J’) Me31B (red), PATR-1 (green) or (K-K’) FMRP (red), PATR-1 (green) and DAPI (blue in I-K and I’-K’). (L-N) Line plots showing fluorescent intensity profiles of (L) TRAL (red) or (M) Me31B (red) or (N) FMRP (red) and PATR-1 (green). The plot shows the normalized pixel-by-pixel gray value of each channel along white lines shown on I’-K’. Computed Pearson’s r values and the statistical significance of the correlation are given above each plot. (O-T) Confocal micrographs of intestinal progenitors from (O) *esg*^*TS*^, (P) *esg*^*TS*^ / *CG18259* RNAi, (Q) *esg*^*TS*^ / *srl* RNAi, (R) *esg*^*TS*^ / *Patr-1* RNAi, (S) *esg*^*TS*^ / *CG13928* RNAi or (T) *esg*^*TS*^ / *me31b* RNAi posterior midguts and stained for TRAL (red), PATR-1 (green) and DAPI (blue). Gray-scale images of TRAL and PATR-1 in panels are shown below the respective panel.

**Figure S3:**
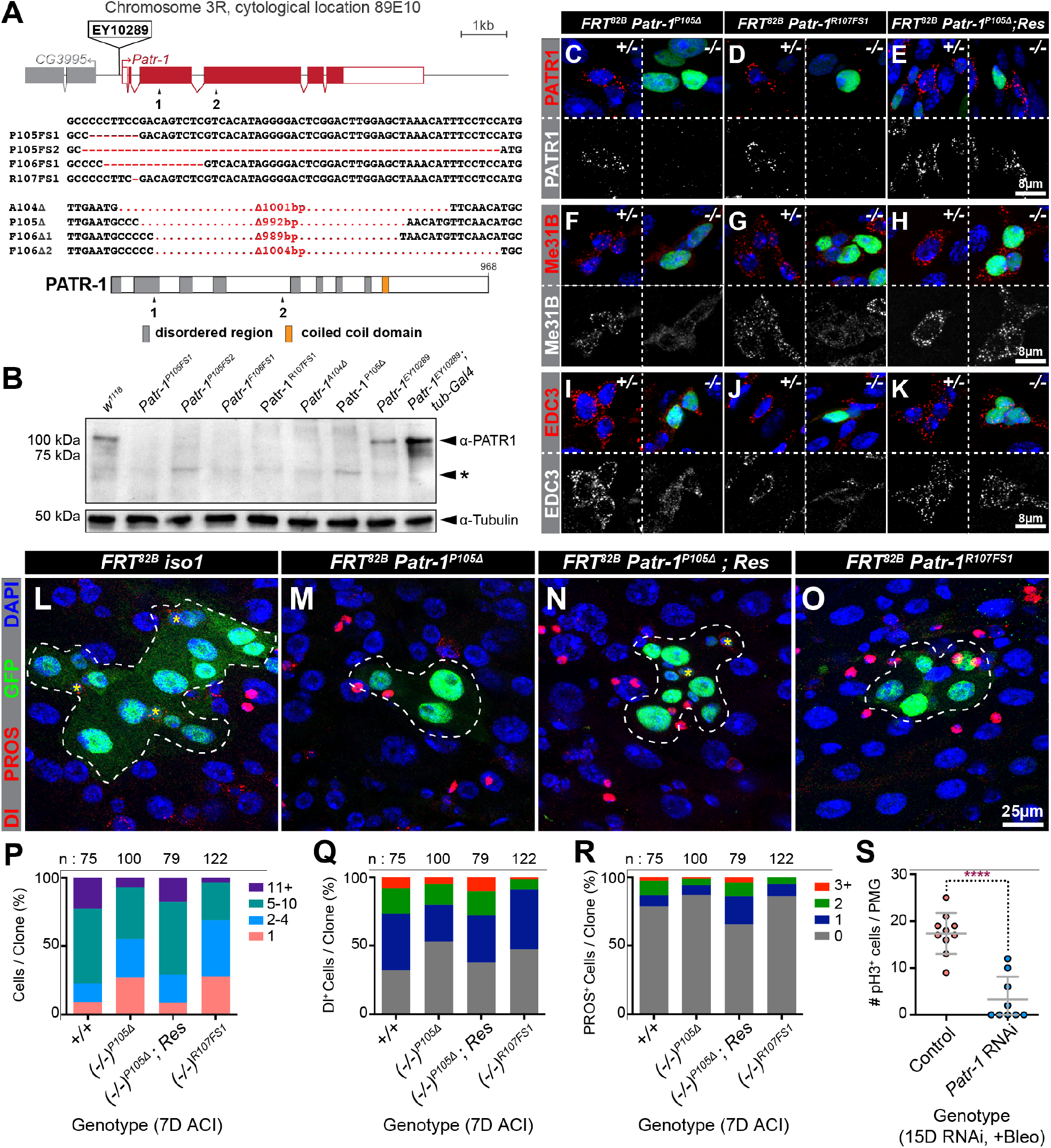
Loss of *Patr-1* blocks P-body formation, reduces the number of ISCs and total number of cells per clone. (A) Schematics of the *Patr-1* locus and PATR-1 protein showing the locations of the two gRNAs (arrowheads) used to generate new alleles as well as the nucleotide changes associated with these alleles. (B) Western blot of *w*^*1118*^ (control), *Patr-1* mutant, and *Patr1*^*EY10289*^ / *tub-GAL4* larval extracts probed with anti-PATR-1 (top panel) or Tubulin (bottom panel) antibodies. A prominent background band detected by the PATR-1 antibody is labeled (*). (C-K) MARCM-labeled homozygous mutant (right, −/−) and neighboring unlabeled heterozygous (left, +/-) *Patr-1*^*P105Δ*^ (C, F, I), *Patr-1*^*R107FS1*^ (D, G, J), or rescued *Patr-1*^*P105Δ*^ (E, H, K) progenitor cells stained for GFP (green), DAPI (blue) and either PATR-1 (red in C-E), Me31B (red in F-H) or EDC3 (red in I-K). (L-O) Representative confocal images of *tub-Gal4, UAS-GFP*-labeled (L) control, (M) *Patr-1*^*P105Δ*^, (N) rescued *Patr-1*^*P105Δ*^, or (O) *Patr-1*^*R107FS1*^ homozygous clones stained for Dl (red), PROS (red), GFP (green) and DAPI (blue). The size of the clone is marked by the white dotted line and ISCs in clones are denoted by the yellow asterisks (*). (P-R) Binned bar plots showing the quantification of (P) total cell, (Q) Dl+ ISC or (R) PROS+ EE cell numbers in *tub-Gal4, UAS-GFP*-labeled control, *Patr-1*^*P105Δ*^, rescued *Patr-1*^*P105Δ*^, or *Patr-1*^*R107FS1*^ homozygous clones analyzed 7 days after clone induction (ACI). Note that n indicates the total number of clones analyzed for each genotype. (S) Dot-plot showing the total number of pH3+ mitotic cells per posterior midgut of *esg*^*TS*^ (n=10) or *esg*^*TS*^ / *Patr-1* RNAi (n=9) flies fed with bleomycin for 24 hours. Rescued (Res) strains harbor the *Patr-1* rescuing transgene, *P21M20*.

**Figure S4:**
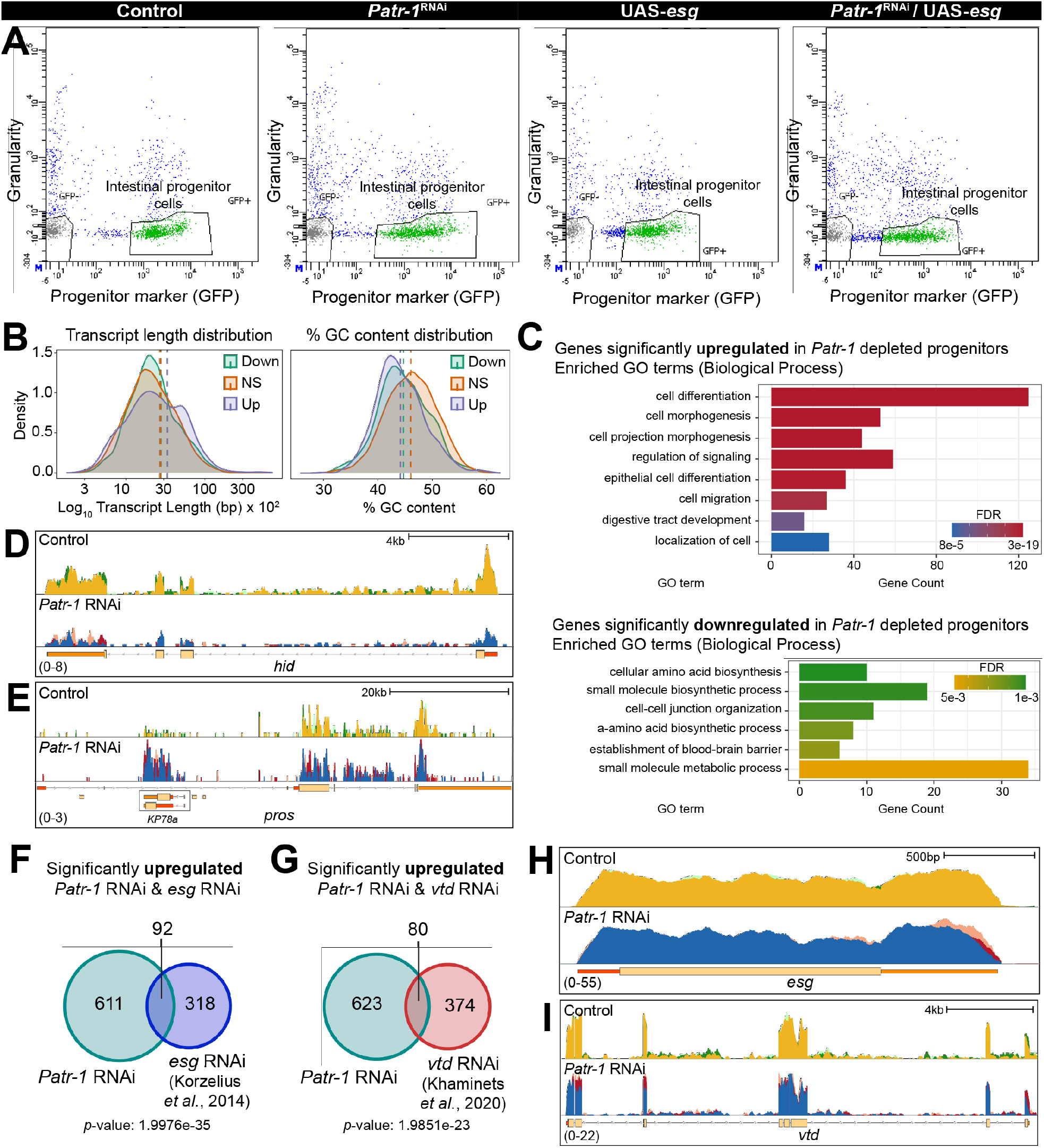
Gene expression changes associated with *Patr-1* RNAi. (A) FACS profiles of *esg*^*TS*^, *esg*^*TS*^ */ Patr-1 RNAi, esg*^*TS*^ */ UAS-esg*, and *esg*^*TS*^ */ Patr-1 RNAi* / *UAS-esg* intestines. (B) Density plot showing the log_10_ transcript length distribution or % GC content of significantly upregulated (Up, purple), downregulated (Down, green) or unchanged (NS, orange) genes. (C) Bar plot showing a selected set of significantly enriched Gene Ontology (GO) terms for genes that are significantly upregulated or downregulated in *Patr-1* RNAi compared to control. (D-E) Genome browser tracks of CPM-normalized control and *Patr-1* RNAi at the (D) *hid* and (G) *pros* loci. (F, G) Venn diagrams of overlap between significantly upregulated genes in *Patr-1* RNAi and (F) *esg* RNAi (Korzelius et al., 2014), or (G) *vtd* RNAi (Khaminets et al., 2020). Note that datasets from Korzelius et al., 2014 and Khaminets et al., 2020 were reanalyzed with *Patr-1* RNAi using the in-house analysis pipeline. (H-I) Genome browser tracks of CPM-normalized control and *Patr-1* RNAi at the (H) *esg* and (I) *vtd* loci. Note the three replicates of the control (yellow, light green and dark green) and the three replicates of the *Patr-1* RNAi (blue, pink and red) on D-E and H-I, have been overlayed for simplicity.

**Figure S5:**
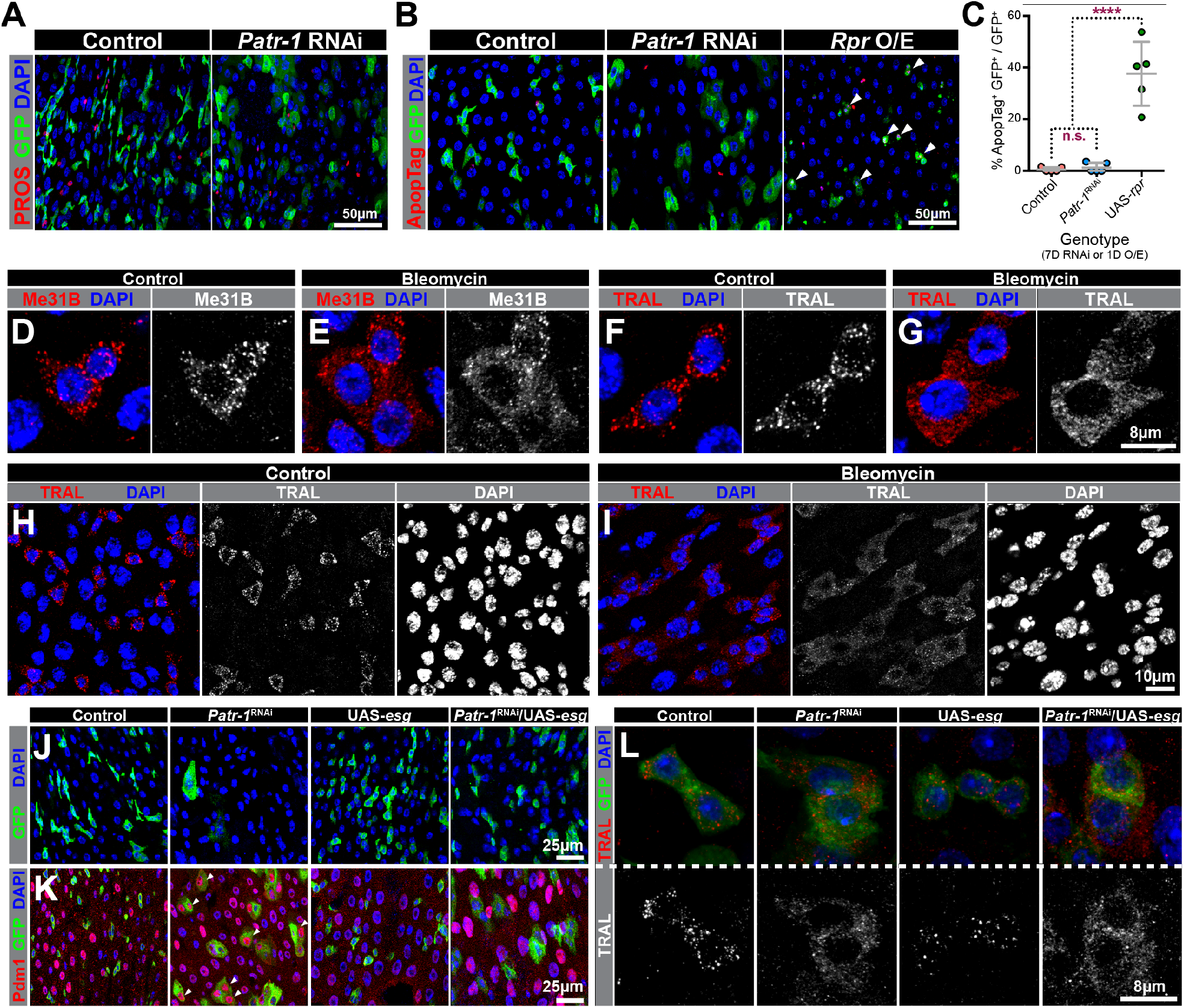
*Patr-1* RNAi does not lead to an increase in the number of EEs or the amount of cell death. (A-B) Confocal micrographs of intestinal sections from (A) *esg*^*TS*^ and *esg*^*TS*^ */ Patr-1* RNAi or (B) *esg*^*TS*^, *esg*^*TS*^ */ Patr-1* RNAi and *esg*^*TS*^ */* UAS-*rpr* and stained for (A) PROS (red) or (B) ApopTag (red) and GFP (green) and DAPI (blue). White arrowheads on B show ApopTag^+^ GFP^+^ cells in the field. (C) Plot of normalized ApopTag^+^ progenitor cells in *esg*^*TS*^ (n=5), *esg*^*TS*^ */ Patr-1* RNAi (n=5), and *esg*^*TS*^ */ UAS-rpr* (n=5) midguts after 7 days of RNAi or 1 day of *rpr* overexpression at 29°C. (D-I) Fluorescent micrographs of intestinal sections from w1118 flies fed with (D, F, H) 5% sucrose in water (control) or (E, G, I) 5% sucrose and 25 µg/ml bleomycin in water (bleomycin) and stained for Me31B (red in D-E) or TRAL (red in F-I) and DAPI (blue in D-I). Note that H-I show broad view images of intestines after each feeding. (J-K) Sections from *esg*^*TS*^, *esg*^*TS*^ */ Patr-1 RNAi, esg*^*TS*^ */ UAS-esg*, or *esg*^*TS*^ */ Patr-1 RNAi* / *UAS-esg* intestines stained for Pdm1 (red, only in K), GFP (green, J-K) and DAPI (blue, J-K). (L) Confocal micrographs of intestinal progenitors from *esg*^*TS*^, *esg*^*TS*^ */ Patr-1* RNAi, *esg*^*TS*^ */ UAS-esg*, or *esg*^*TS*^ */ Patr-1 RNAi / UAS-esg* posterior midguts stained for TRAL (red), GFP (green) and DAPI (blue). Error bars on plots show mean±s.d. and asterisks denote statistical significance from Ordinary one-way ANOVA with Turkey’s multiple comparison test (H). \*\*\*\**p* < 0.0001, n.s. = not significant.

